# *Schistosoma mansoni* infection reprograms the metabolic potential of the myeloid lineage in a mouse model of metabolic syndrome

**DOI:** 10.1101/2020.04.20.050898

**Authors:** Diana Cortes-Selva, Lisa Gibbs, J. Alan Maschek, Tyler Van Ry, Bartek Rajwa, James E. Cox, Eyal Amiel, Keke C. Fairfax

**Affiliations:** Department of Pathology, Division of Microbiology and Immunology, University of Utah, Salt Lake City UT, 84112; Bindley Bioscience Center, Purdue University, West Lafayette IN, 47907; Department of Biochemistry, University of Utah, Salt Lake City UT, 84112; Department of Biomedical and Health Sciences, University of Vermont, Burlington, VT 05405; Metabolomics, Proteomics and Mass Spectrometry Cores, University of Utah, Salt Lake City, UT, 84112; Department of Nutrition and Integrative Physiology and the Diabetes and Metabolism Research Center, University of Utah, Salt Lake City, UT 84112

**Keywords:** Myeloid lineage, macrophage metabolism, *Schistosoma mansoni*, biological sex, metabolic disease

## Abstract

Despite evidence that helminths protect from metabolic disease, a major gap exists in understanding the underlying mechanism(s). Here we demonstrate that bone marrow derived macrophages (BMDM) from *S. mansoni* infected male ApoE^-/-^ mice have dramatically increased mitochondrial respiration compared to those from uninfected mice. This change associates with increased glucose and palmitate shuttling into TCA cycle intermediates and decreased accumulation of cellular cholesterol esters. Moreover, systemic metabolic modulation by schistosomes is a function of biological sex, where infection protects ApoE^-/-^ male, but not female, mice from obesity and glucose intolerance. Sex-dependence extends to myeloid cells, where reprogramming leads to opposite cholesterol phenotypes in BMDM from females and males. Finally, the metabolic reprogramming of male myeloid cells is transferrable via bone marrow transplantation to an uninfected host, indicating maintenance of reprogramming in the absence of sustained antigen exposure. This work reveals that *S. mansoni* systemic reprograming of myeloid metabolism is sex-dependent.

## Introduction

Cardiovascular disease (CVD) is the leading worldwide cause of mortality (Hinton et al., 2018; Roth et al., 2017). In the United States, 65% of adults diagnosed with diabetes have elevated LDL cholesterol levels or take cholesterol lowering medications, and death rates from atherosclerotic cardiovascular disease (CVD) are ∼1.7 times higher in this population as compared to non-diabetic adults (Emerging Risk Factors et al., 2010). It is well established that in the diabetic population, obesity, and dyslipidemia are risk factors underlying these increases in mortality, while hyperglycemia is an independent risk factor (Marks and Raskin, 2000; Wong et al., 2016). Underlying conditions such as diabetes and atherosclerosis contribute to the burden of CVD in both females and males. While, the incidence of CVD is markedly higher in men than in age-matched women (Opotowsky et al., 2007; Tan et al., 2010), the risk of developing CVD while diabetic is much greater in women than men (Humphries et al., 2017; Peters et al., 2014). In non-diabetic patients, females exhibit increased insulin sensitivity in comparison to males, as well as reduced prevalence of dysglycemia and enhanced muscle glucose uptake (Cnop et al., 2003; Kim and Reaven, 2013; Moran et al., 2008; Willeit et al., 1997), suggesting sex-dependent modulations in whole body metabolism. Recent studies suggest a role of the gut microbiota in the differences between sexes in the regulation of lipid metabolism (Baars et al., 2018). Nevertheless, complete understanding of mechanistic basis behind sexual dimorphism in metabolic syndrome is still lacking.

Previous studies have uncovered an association between a history of helminth infection and reduced prevalence of metabolic disease in humans and rodents ((Doenhoff et al., 2002; Stanley et al., 2009; Wiria et al., 2015). Specifically, infection by Schistosomes reduces cholesterol and atherosclerotic plaques (Doenhoff et al., 2002; Stanley et al., 2009), this effect has been attributed, in part, to an anti-inflammatory phenotype in macrophages (Wolfs et al., 2014) and transcriptional reprogramming of phospholipid and glucose metabolism related genes in hepatic macrophages (Cortes-Selva et al., 2018). Moreover, it has been postulated that schistosomes have the potential to affect long term glucose metabolism in T cells (Chen et al., 2013). Accumulating evidence suggests that biological sex affects disease progression; yet the effect of schistosomiasis on metabolic-protection in females and males it is not well understood, as most studies have been conducted only in males or no sex differentiation has been made during data analysis ((Sanya et al., 2019; Shen et al., 2015; Wolde et al., 2019)). To date, no report with a particular emphasis on females has been conducted.

Schistosomiasis induces Th2 polarization and alternative activation of macrophages, essential for host survival (Barron and Wynn, 2011; Fairfax et al., 2012; Herbert et al., 2004). IL-4 induced alternative activation of macrophages relies on oxidative phosphorylation (OXPHOS) and fatty acid oxidation for energy production, and is dependent on cell intrinsic lysosomal lipolysis (Huang et al., 2014; Vats et al., 2006). Macrophage metabolism follows a dysmorphic pattern, as sex-related differences affect the processes involved in cholesterol and lipid metabolism in macrophages as well as inflammatory cytokine production in adipose tissue (Griffin et al., 2016; Ng et al., 2001). Moreover, in rats, phagocytes from females had increased ROS generation than males (Rudyk et al., 2018). Such differences have often been attributed to the role of sex hormones in gene expression and immune cell function (Rubinow, 2018; Taneja, 2018; Winn et al., 2019), but a clear understanding of the effects of sex on the regulation of macrophage metabolism, as well as how sex modulates the effects of schistosomiasis in the protection from metabolic disease is lacking.

In the present study, we sought to determine the systemic effects of *S. mansoni* infection on the myeloid lineage. Surprisingly, we discovered that macrophages derived from the bone marrow of *S. mansoni* infected male mice have dramatically increased oxygen consumption and mitochondrial mass compared to those from uninfected males. This shift is accompanied by increased carbon shuttling into TCA cycle intermediates, a decrease in cholesterol esters, and increased fatty acid oxidation. When we examined the role of biological sex in schistosome induced modulation we found that *S. mansoni* infection does not reliably protect ApoE^-/-^ female mice from HFD induced weight gain or glucose intolerance. The sex-dependent effect of infection extends to the myeloid lineage, where bone marrow derived macrophages from infected females display the opposite metabolic phenotype as those from infected males, with a dramatic increase in cellular cholesterol esters. Overall, these data present the first evidence that *S. mansoni* systemically modulates the myeloid compartment in a sex-dependent manner and provide a more comprehensive understanding of how *S. mansoni* infection may confer metabolic protection at the cellular level.

## Results

### Macrophages derived from *S. mansoni* infected male mice have increased oxygen consumption and spare respiratory capacity

We have previously reported that schistosomiasis alters the expression of numerous genes relevant to glucose, cholesterol, and amino acid metabolism in hepatic macrophages of male mice (Cortes-Selva et al., 2018). These alterations are associated with improved insulin sensitivity and atherosclerotic score in male mice. Since it has previously been shown that during *S. mansoni* infection the majority of liver macrophages are monocyte derived, and monocyte recruitment drives both atherosclerosis (Potteaux et al., 2011; Tacke et al., 2007) and obesity induced insulin resistance (Beliard et al., 2017; Liang et al., 2007; Oh et al., 2012; Rull et al., 2010), we hypothesized that Schistosome infection may imprint the monocyte-macrophage lineage with altered metabolic propensities. To elucidate whether infection imprints macrophages with an altered metabolic phenotype we infected (and mock infected controls) atherogenesis-prone male ApoE^-/-^mice on high-fat diet (HFD), and sacrificed them at 10-weeks post-infection to harvest bone marrow cells. Macrophages were differentiated *in vitro* with M-CSF in a 6-7-day culture. We performed real-time extracellular flux analysis on unstimulated bone marrow derived macrophages (BMDM) from ApoE^-/-^ HFD infected and uninfected (control) HFD mice to quantify oxygen consumption rate (OCR) (Figure 1A). BMDM from ApoE^-/-^ HFD infected mice showed improved basal respiration (Figure 1B) and significantly increased spare respiratory capacity (p<0.0001, Figure 1C). Since eukaryotic cells integrate oxidative phosphorylation (OXPHOS), glycolysis and the tricarboxylic acid (TCA) cycle to satisfy energy requirements, we also tested the extracellular acidification rate (ECAR), which is suggested as a marker of inhibited mitochondrial respiration (Pike Winer and Wu, 2014), in BMDM from infected and uninfected ApoE^-/-^ HFD mice. We observed no differences in ECAR in infected male mice compared to uninfected controls (Figure 1D). Cell intrinsic lysosomal lipolysis has previously been shown to support macrophage spare respiratory capacity in the context of macrophage alternative activation (Huang et al., 2014; Liu et al., 2017). We stained for hydrophobic and neutral lipids by Oil Red O (ORO) (Mehlem et al., 2013) and observed that the lipid content of BMDM from infected ApoE^-/-^ males trended to reduction in comparison to the control group, but was not significantly reduced (Figure 1E). To analyze BMDM mitochondrial mass, which has also been linked to increased respiratory capacity (Langston et al., 2017), we analyzed mitochondrial activity by Mitotracker Deep Red FM. We observed that BMDM from infected mice exhibited increased MitoTracker median fluorescent intensity (MFI) in comparison to the BMDM from uninfected mice (Figure 1F). Overall, these data indicate that *S. mansoni* infection in males leads to increased oxygen consumption and mitochondrial metabolism in BMDM. Mitochondrial oxidative dysfunction in macrophages has recently been linked to insulin resistance (Jung et al., 2018), so this metabolic shift could contribute to the infection-induced improvement in glucose tolerance seen in infected males.

**Figure 1.**
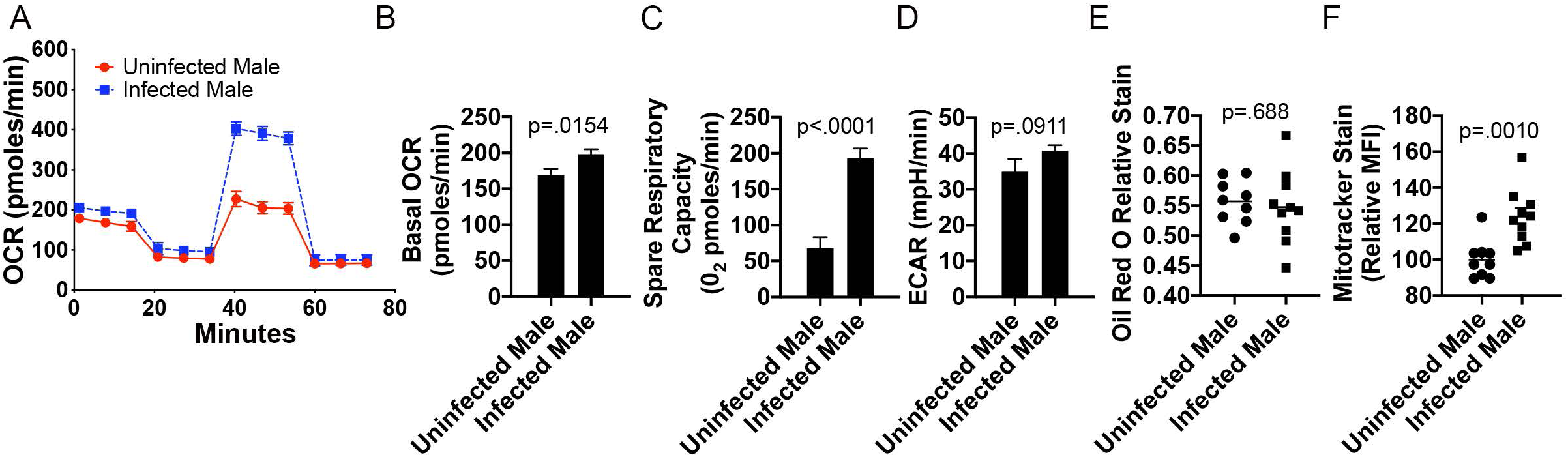
Bone marrow derived macrophages (BMDM) from ApoE^-/-^ male *S. mansoni* infected mice exhibit increased oxygen consumption and mitochondria mass. ApoE^-/-^ male were fed HFD for 10 days before infection with *S. mansoni*. Ten weeks post infection mice were sacrificed and bone marrow cells were harvested and cultured for 7 days under M-CSF. (A) SeaHorse assay results for OCR of BMDC from infected and uninfected ApoE^-/-^ males in basal conditions and in response to mitochondrial inhibitors. (B) Quantification (in picomoles per minute) of the basal oxygen consumption of BMDM from uninfected or infected ApoE^-/-^ HFD male mice. (C) Quantification of the spare respiratory capacity of BMDM from uninfected or infected ApoE^-/-^ HFD male mice (D) Extracellular acidification rate of BMDM from male uninfected or infected ApoE^-/-^. (E) Oil Red O relative staining in BMDM from ApoE^-/-^ mice (F) MitoTracker Red Deep Stain measure by flow cytometry in BMDM from ApoE^-/-^ mice. SeaHorse assay analysis were performed the Seahorse XFe96 instrument. *p < 0.05; **p < 0.01; ***p < 0.001. Graphs are representative of multiple experiments (2-3), with n>4 per group.

### Schistosomiasis in male ApoE^-/-^ mice alters metabolic flux of glucose and the lipidomic fingerprint of macrophages

In order to understand how Schistosome infection alters the metabolic fingerprint and promotes mitochondrial metabolism in macrophages derived from ApoE^-/-^ HFD male mice, we performed metabolic tracing analysis, where macrophages were differentiated in the presence of normal glucose and then switched to ^13^C-labeled glucose for 24 hours. We observed increased shuttling of heavy labeled glucose to malate (Figure 2A), citrate (Figure 2B), itaconate (Figure 2C), and succinate Figure 2E in BMDM from infected mice in comparison to BMDM from uninfected mice. The lack of increased heavy lactate production (Figure 2D), suggests that the primary reprogramming is focused on glucose-dependent mitochondrial metabolism. Importantly, we found no upregulation of alternative activation markers (CD301, CD206, Arg1, Nos2) in unstimulated BMDM from infected mice when compared to BMDM from controls (Supplementary Figure 1). Since we have previously shown that infection alters the phospholipid and cholesterol metabolism in hepatic macrophages (Cortes-Selva et al., 2018), we performed unbiased lipidomics using liquid chromatography-mass spectrometry (LC-MS). We used a supervised model by partial least squares-discriminant analysis (PLS-DA), with two components to determine the lipidomic profile of BMDM. There was a robust separation between groups that indicates the metabolic profiles of BMDM from infected and uninfected males differ significantly and suggests that there is a prominent alteration of metabolites induced by infection in male mice (Figure 2F). The lipid species that drive the variation observed in the PLS-DA as measured by the Variable Importance in Projection (VIP) score from uninfected males in comparison to infected males included cholesterol esters (CE) (20:1), CE (22:0), CE (24:0), CE (24:1), CE (22:4), CE (22:1), CE (18:0); diacylglycerols (DG) (16:0_16:0), DG (16:0_18:0) and triacyclglycerol (TG) (16:0_16:0_16:0) (Figure 2G,H). We then analyzed the total CE levels in macrophages from both groups of male mice and found that infection led to significantly reduced CE in male mice (p<0.0001, Figure 2I), further evidencing an important role of cholesterol metabolism in macrophages following helminth infection. Next, we determined the transcriptomic modifications induced by infection in unstimulated BMDM from infected and uninfected males by mRNA sequencing (mRNAseq). Significant gene expression differences were observed in BMDM from infected male mice, compared to uninfected controls. Transcripts from the two groups are depicted in Volcano plots, using false discovery rate (FDR<0.05 in red, FDR<0.01 in blue) and Log2 fold changes (cut off of .6 Log2 FC) to identify statistically significant genes (Figure 2J). Among the differentially regulated factors are Gbp6, which is related to interferon-γ signaling and innate immune function; Gm7609, a predicted pseudogene; Gbp4, a member of the GTPase family involved in protective immunity against microbial and viral pathogens; Apol9a, which is predicted to be related to lipid transport, lipoprotein metabolic processes and stimulated by interferon; Iigp1, a GTPase with roles in response to intracellular pathogens; CD300e, which belongs to families of paired activating and inhibitory receptors implicated in immune responses; and Batf2, a basic leucine zipper factor whose activation is detrimental for type-1 infectious disease. Importantly, Mgll, which encodes monoacylglycerol lipase that catalyzes the conversion of monoacylglycerides to free fatty acids and glycerol was significantly increased. Mgll is required for lipolysis and improved glucose homeostasis in mice on HFD (Berdan et al., 2016; Douglass et al., 2015), an infection driven increase was validated by RT-qPCR (Figure 2L). In addition, Slc1a3, which encodes for the glutamate aspartate transporter 1 that is localized in the inner mitochondria membrane as part of the malate-aspartate shuttle and is relevant for amino acid homeostasis in adipocytes, was significantly altered in our model, and was further validated by RT-qPCR (Figure 2M). Interestingly, the pathways significantly altered (Figure 2K) during infection included hematopoietic cell lineage (p=5.779 ×10^−8^), asthma (p=3.169 ×10^−5^), and cytokine-cytokine receptor interactions, graft-versus host disease, type 1 diabetes and allograft regression (p=0.0000894). Together, these data identify key factors in immune and metabolic responses as well as novel factors with unknown functions that are regulated by exposure to *S. mansoni* infection.

**Figure 2.**
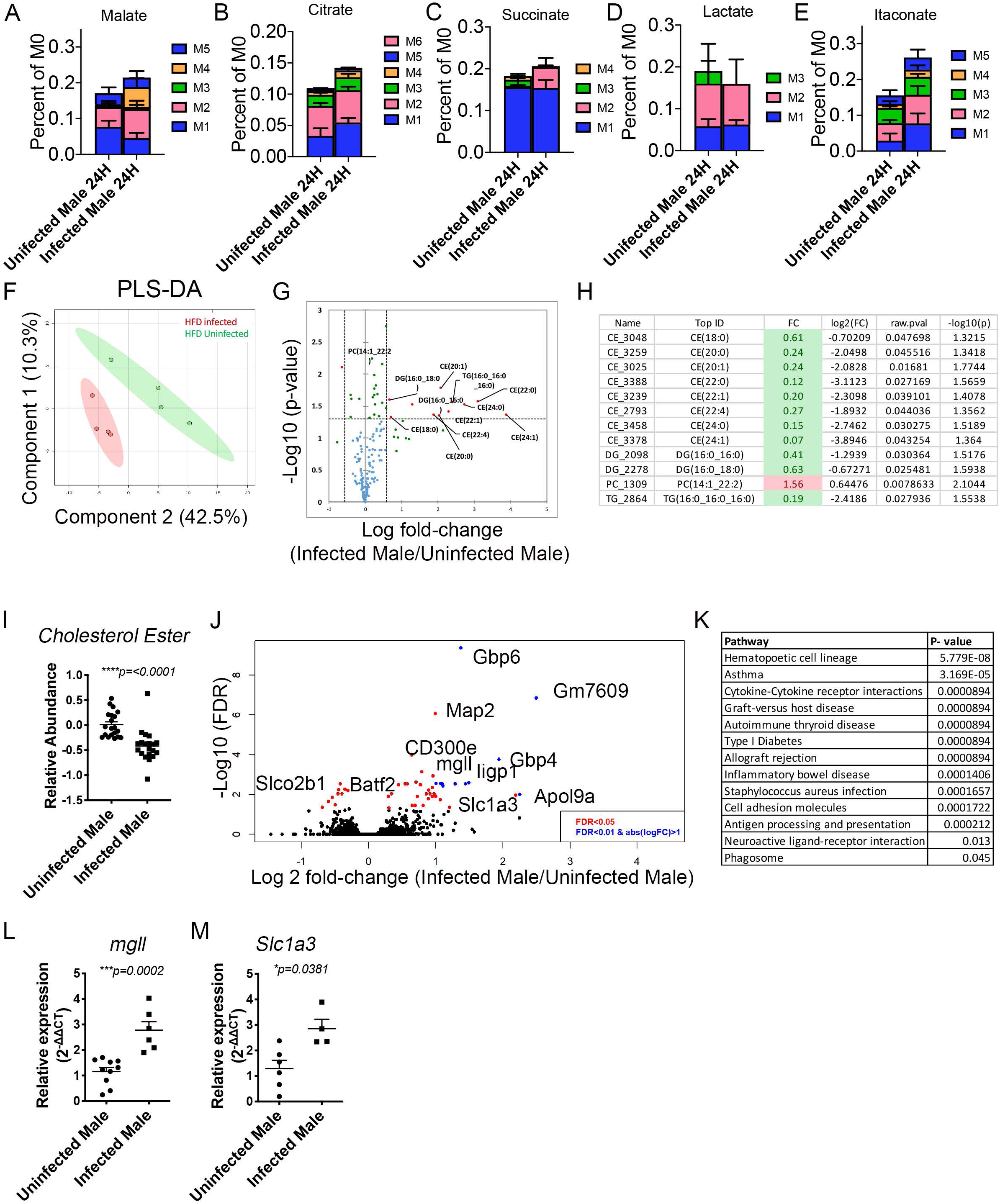
BMDM from male *S. mansoni* infected HFD ApoE^-/-^ mice have increased TCA cycle usage and significantly reduced cholesterol esters. A-E) Mϕ were differentiated from bone marrow of 10-week infected animals with M-CSF in a 7-day culture in normal glucose and then switched to ^13^C-labeled glucose for 24 hours. F-H) Mϕs were differentiated with M-CSF in a 7-day culture and then total cellular lipids were extracted and analyzed via LC-MS based lipidomic analysis. F) PLS-DA derived score from HFD Infected and HFD Uninfected BMDM. G) Plot of lipid species from the BMDM of infected and uninfected males on HFD, significantly altered lipid species are red and labeled. H)Table of statistical analysis of VIP I) Box whisker plot of normalized AUC of cholesterol ester species, which were identified as VIP compounds from the PLS-DA analysis. J-L) Bone marrow macrophages were differentiated with M-CSF and mRNA sequenced via RNASeq. J) Volcano plot of significantly differentially expressed genes between BMDM from *S. mansoni* infected uninfected mice. K) iPathway analysis showed distinct profiles in BMDM from *S. mansoni* infected, mice. L,M) Total RNA was extracted from biologically independent BMDM differentiated in M-CSF for Real-time PCR validation of mgll and slc1a3 regulation. Data in A-I are representative of 3 experiments with 4-6 mice per group in each experiment. Data in J and K are representative of sequencing of 2 biologically independent experiments with 5-6 mice per group. Data in L are from 2 biologically independent experiments with 4-6 mice per group.

### *S. mansoni* infection protects male mice, but not female mice, from obesity and glucose intolerance independently of systemic alternative macrophage activation

Sex is a key contributor to the phenotype of cardiovascular and metabolic features in mammals (Chella Krishnan et al., 2018), so we assessed the sex dependent impact of *S. mansoni* infection on obesity and glucose intolerance. For this, we fed male and female ApoE^-/-^ HFD for 10-days before infection. We infected and mock infected mice (controls) as described in the methods. Ten weeks post infection we analyzed body weight and glucose tolerance (via an IP glucose tolerance test) and found that infection is significantly beneficial for male, but not female mice, as only males are protected from HFD-induced weight gain and glucose intolerance (Figure 3A, 3B). We also analyzed serum triglyceride (TG) and diglyceride (DG) levels using untargeted lipidomics and found that relative abundance of both TG and DG were decreased in infected males as compared to uninfected males, while TGs were increased by infection in females and DGs were unchanged (Figure 3C). These data indicate that *S. mansoni* infection induces a sex-dependent modulation of metabolic disease parameters. We then wondered if this infection-mediated effect in males only was correlated with differences in systemic alternative activation of hepatic macrophages in females. Flow cytometry analysis showed that alternative activation markers (CD206, CD301, Arg1) were highly expressed in macrophages from male and female mice following infection, but not in naïve mice (Figure 3D), suggesting that *S. mansoni* induced alternative activation irrespective of sex. Previous studies characterizing the dynamics of alternatively activated macrophages during schistosome infection have found that these macrophages primarily arise from Ly6C^high^ monocytes (Girgis et al., 2014; Nascimento et al., 2014). Naïve male and female ApoE^-/-^ mice on HFD had equivalent frequencies of Ly6c^high^ monocytes circulating in peripheral blood. At 10-weeks post infection we found an increased percentage of both Ly6C^int^ and Ly6C^high^ cells (Figure 3E) in both male and female mice compared to the mock infected controls, with the frequency of Ly6C^high^ cells in females 1.74 times that of males, suggesting either increased monopoiesis in females, or increased tissue recruitment in males. Since we had found increased mitochondrial MFI in BMDM from infected male mice, we asked whether circulating blood monocytes are similarly modulated. We observed an infection induced increase in Mitotracker fluorescent intensity in monocytes from male ApoE^-/-^ mice on HFD, but not from females (Figure 3F). These data suggest that there is sex-specific increased mitochondrial activity following infection in the monocyte cell population. We then wondered if these effects in the differentiated monocytes are the result of long-lasting changes in the myeloid lineage after helminth infection. For this, we analyzed the main lineages of hematopoietic progenitors that produce myeloid cells: granulocyte-monocyte progenitors (GMP), monocyte-DC progenitors (MDP) and the common myeloid progenitor (CMP) in female and male ApoE^-/-^ mice. CMP were defined as Lin^-^CD127^-^c-Kit^+^Sca-1^-^CD34^+^FcRII/III^lo/-^(Paul et al., 2016), GMP were defined as Lin^-^IL-7R^-^Sca-1^-^c-kit^+^CD34^+^FcR II/III^+^, and MDP were defined as defined as Lin^-^ c-Kit^+^ Sca-1^-^ CD34^+^ FcγR^lo^ CD115^hi^ cells. Overall bone marrow cellularity was not affected by Schistosome infection (Figure 3G), however, the numbers of CMP and GMP in infected male mice were significantly reduced compared to uninfected controls, while GMP and CMP were not reduced in females (Figure 3 H-J). The numbers and percentages of MDP remained unchanged in infected females and males compared to uninfected controls (Figure 3K, 3L). These data suggest that *S. mansoni* infection modulates both metabolic disease and the myeloid lineage in a sex-specific manner that is independent from the induction of systemic alternative activation.

**Figure 3.**
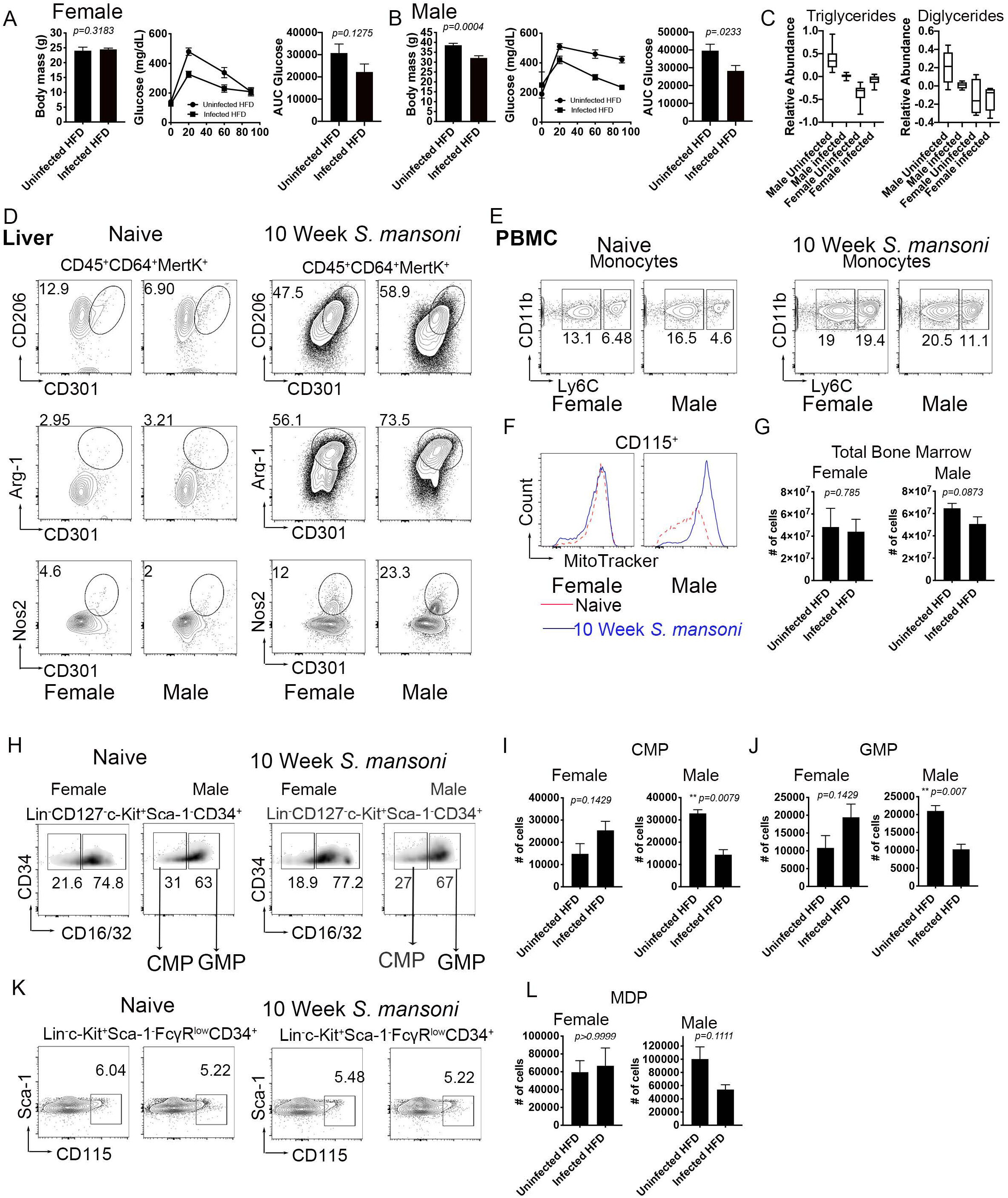
*S. mansoni* infection does not protect females from metabolic disease despite inducing alternative activation in hepatic macrophages. (A-B) Total body weight and glucose tolerance test (GTT) at 10 weeks post-infection. GTT values were analyzed by Area Under the Curve and graphed using GraphPad Prims. (C) (D) Flow cytometry analysis of alternative activation markers CD206 and CD302 in perfused and digested livers gated on hepatic macrophages (CD45^+^CD64^+^MertK^+^) Arg-1 expression in hepatic macrophages from uninfected and infected ApoE^-/-^ mice Nos-2 expression by flow cytometry in hepatic macrophages 10 weeks post infection. (E) Ly6C expression in monocyte from peripheral blood mononuclear cells (PBMC) at 10-week post infection by flow cytometry (F) MitoTracker Red in CD115^+^ monocytes from PBMC at 10 weeks post immunization in uninfected and infected ApoE^-/-^ mice. (G) Total cell counts from bone marrow cells utilizing trypan-blue discrimination of apoptotic cells (H) Percentages of CMP (Lin^-^CD127^-^c-Kit^+^Sca-1^-^CD34^+^CD16/32^low^) and GMP (Lin^-^CD127^-^c-Kit^+^Sca-1^-^CD34^+^CD16/32^hi^) (I) CMP cell counts in bone marrow ApoE^-/-^ (J) GMP cell counts 10 week post infection in ApoE^-/-^ mice (K) Flow cytometry analysis of MDP defined as Lin^-^c-Kit^+^Sca-1^-^FcγR^low^CD34^+^CD115^+^ (L) MDP cell counts in ApoE^-/-^uninfected and infected male mice. Graphs are representative from experiments that were performed 3-4 times with n>4. Exception is MitoTracker data, which was performed twice, with n>4. Statistical analysis was done using unpaired Student’s t test, *p < 0.05; **p < 0.01.

### *S. mansoni*-increases fatty acid oxidation in male but not female ApoE^-/-^ mice on HFD

In order to determine whether infection-induced pathologic differences in males and females were accompanied by sex-specific differential macrophage metabolic regulation, we cultured bone marrow cells from 10-week infected or uninfected control male and female mice for 7 days to generate BMDM. OCR was measured in real time in basal conditions and following the addition of mitochondrial inhibitors in unstimulated BMDM from both infected and uninfected male and female ApoE^-/-^ animals. Similar to as seen in Figure 1A, BMDM from infected males exhibited increased basal OCR and spare respiratory capacity compared to uninfected male controls (Figure 4A, 4B, 4C). However, basal OCR and spare respiratory capacity of BMDM from infected females remained unaltered in comparison to BMDM from uninfected females (Figure 4A, 4B, 4C). Moreover, side by side OCR analysis in females and males with palmitate as a substrate (glucose limiting conditions) showed that BMDM from infected male, but not from infected females had an increased ability to oxidize exogenous palmitate (Figure 4D). In addition, BMDM from infected males but not females had significantly increased palmitate basal OCR and palmitate spare respiratory capacity, suggesting exogenous free fatty usage as a carbon source for OXPHOS (Figure 4E, 4F). Similar to the male only data, we observed no differences in macrophage lipid content in either group, suggesting that global lipolysis may not underlie OCR and spare respiratory capacity in our model (Figure 4G). Additionally, analysis of mitochondrial mass in females and males showed that similar to blood monocytes, BMDM from infected male mice, but not females have a significantly higher Mitotracker MFI. These data suggest that *S. mansoni* infection improved mitochondrial biogenesis in males, but not females. Again, suggesting differential regulation of macrophage metabolism based on biological sex. This regulation may account in part for the differences in the role of infection in modulating obesity and insulin sensitivity.

**Figure 4.**
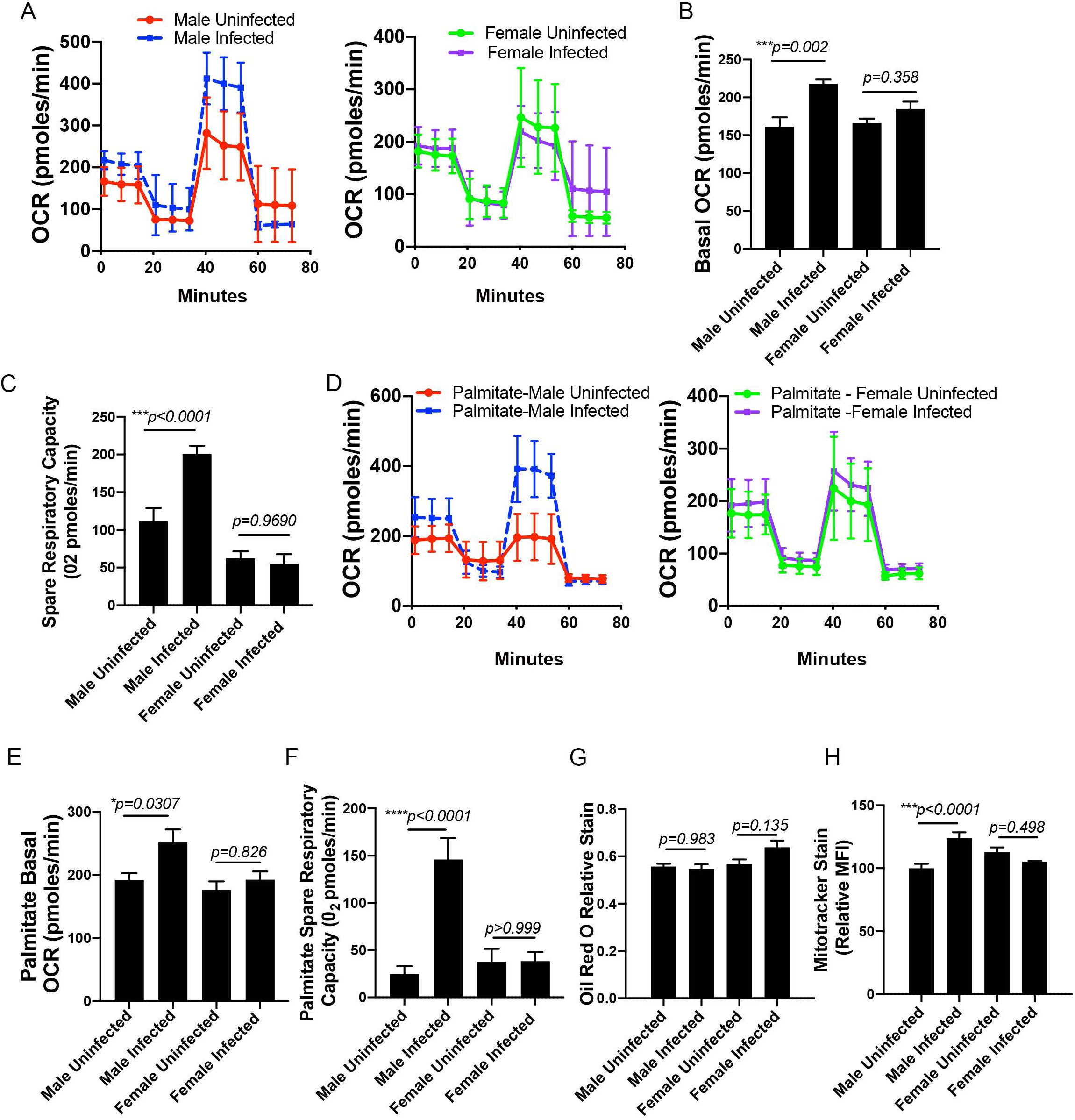
*S. mansoni* infection induces sex-specific modulation of oxygen consumption and beta-oxidation in BMDM. (A-C) oxygen consumption rate and spare respiratory capacity were measured at steady state. D-F) BMDM palmitate dependent oxygen consumption and spare respiratory capacity was measured in glucose limiting conditions. G) Oil Red O relative staining in BMDM from ApoE^-/-^ mice (H) MitoTracker Red Deep Stain measure by flow cytometry in BMDM from ApoE^-/-^ mice. SeaHorse assay analysis were performed the Seahorse XFe96 instrument. *p < 0.05; **p < 0.01; ***p < 0.001. Graphs are representative of multiple experiments (2-3), with n>4 per group.

### *S. mansoni* infection differentially alters the cellular lipid profile in female and male mice

To understand the sex specific modulations induced by schistosomiasis in macrophage lipid metabolism, we isolated cellular lipids from unstimulated BMDM and performed unbiased lipidomics in male and female derived-cells. PLS-DA showed that *S. mansoni* infection led to a sex-specific lipid profile (Figure 5A). Further, analysis of total CE (identified as a VIP species in males in Figure 2) showed that infection led to decreased levels of CE in cells derived from infected male mice. In contrast, infection led to significantly increased total CE in BMDM derived from infected female mice (Figure 5B). Similar to males, infection in females induces a unique lipid signature (Figure 5C), with two species of plasmanyl-PE, three species of plasmanyl-PC, BMP and CE as the main drivers of the altered lipid profile in females during infection (Figure 5D). Since we observed BMP’s as a VIP compound driven by *S. mansoni* infection in females, but not males, we went back and performed targeted lipidomics to quantify BMPs at the class level. Indeed, total BMPs were significantly increased by infection in BMDM from females but not males (Figure 5E). Our Seahorse data indicates that *S. mansoni* infection induced differential oxidation of exogenous palmitate in BMDM derived from males and females. In order to understand the metabolic flux of exogenous palmitate, we performed metabolic tracing analysis of palmitate, where macrophages were differentiated in the presence of normal glucose and then switched to C^13^-labeled palmitate for 24 hours. Similar to what we documented with glucose, we observed increased shuttling of heavy labeled palmitate into succinate, malate, and fumarate in BMDM derived from infected males, with decreased shuttling into myristate (a marker of fatty acid elongation). BMDM derived from infected females displayed the opposite phenotype, with decreased shuttling of heavy labeled palmitate into succinate, malate, and fumarate, and increased shuttling into myristate compared to BMDM from uninfected controls. These data suggest that BMDM derived from infected male ApoE^-/-^ mice have increased basal utilization of exogenous palmitate for beta-oxidation, while the female data suggests increased lipid synthesis, strengthening the data obtained from our palmitate extracellular flux analysis.

To further understand the role of biological sex in the *S. mansoni* induced reprograming of the BMDM lipidome, we isolated cellular lipids from male and female BMDM either unstimulated (media), or following LPS stimulation, and then performed untargeted lipidomics as described in Figure 5. We analyzed the lipidomic data with machine learning using a two-step selection process (see Methods). This approach identified ∼20 features (lipid compounds) contributing to an elastic net regression model that can predict the infection status of samples-originating animals (Zou, 2005). These informative features are listed in descending order of importance (Supplemental Figure 2A). Interestingly, some key features (for instance, the top scoring one) are not represented as significant in the univariate tests (univariate adjusted p values, Supplemental Figure 2B). However, these features are important contributors to the multivariate machine learning model. This is not unexpected – some features that would be considered uninformative individually may be very useful in improving predictions if combined with other features. When we examine these features individually, we can see a differential response to infection for males and females (for instance, PG 18:1_18:2, a glycerophospholipid) represented in Supplemental Figure 2C). Seven of the 20 molecular features identified by machine learning are BMP (Bis(monoacylglycerol)phosphate) species, which have been implicated in glycosphingolipid degradation and cholesterol transport (Anheuser et al., 2019; Luquain-Costaz et al., 2013). These data further support the hypothesis that *S. mansoni* infection differentially modulates the metabolism of myeloid cells from male and female animals, and that this modulation revolves around decreased cholesterol storage and fatty acid synthesis in males and increased cholesterol storage and fatty acid synthesis in females. Cholesterol and lipid metabolism has previously been associated with inflammatory myeloid effector function (Carroll et al., 2018; Funk et al., 1993; Oiknine and Aviram, 1992), so we quantified the acute inflammatory effectors nitrite and iNOS, along with pro-inflammatory cytokines/chemokines IL-12p70, CXCL1, and IL-6 (chemokines/cytokines with known pathogenic roles in obesity, insulin resistance and atherosclerosis) following stimulation with LPS. *S. mansoni* infection increases LPS induced nitrite and iNOS production in BMDM from male, but not female ApoE^-/-^ mice on HFD (Figure 5 J,K). Conversely, BMDM from infected males have decreased production of IL-2p70, CXCL1, and IL-6 (Figure 5 L-N) following LPS stimulation as compared to BMDM from uninfected controls. LPS induced IL-12p70, CXCL1, and IL-6 production by female BMDM is unaffected by infection status. Increased production of the effector molecules nitrite and iNOS combined with decreased production of pro-inflammatory mediators associated with chronic inflammation supports the idea that *S. mansoni* infection promotes a hybrid macrophage state in males.

**Figure 5.**
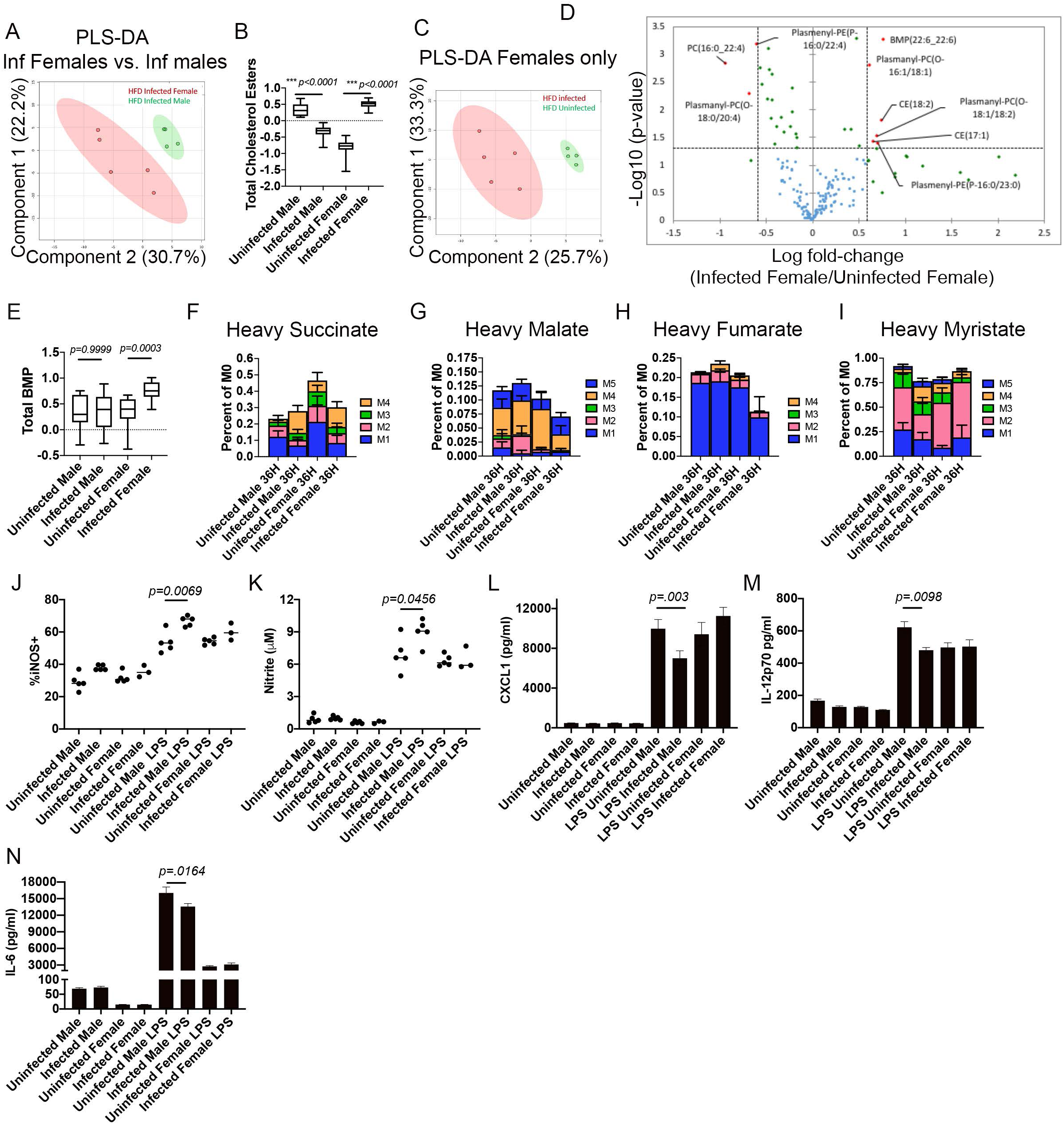
*S. mansoni* infection induces significantly different cellular lipid profiles in BMDM from HFD female and male ApoE^-/-^ mice. A) PLS-DA derived score of LC-MS based lipidomic analysis of BMDM from HFD Infected females vs males. B) Box whisker plot of normalized AUC of cholesterol ester species, between female and male infected and uninfected HFD ApoE^-/-^ mice. C) PLS-DA derived score from Female HFD Infected and uninfected BMDM. D) Dot plot of lipid species identified in BMDM from infected and uninfected female ApoE^-/-^ mice on HFD, significant lipid species are represented by red dots and labeled. E) Relative quantitation of BMP compounds in BMDM from male and female animals with and without *S. man*soni infection. F-I) Macrophages were differentiated with M-CSF in a 7-day culture with normal glucose and serum, and then switched to ^13^C-labeled palmitate in dialyzed serum for 36 hours. J-N) Culture supernatants from BMDM stimulated with LPS for 24 hours were assayed for chemokines/cytokines. Data in A-E and J-N are representative of 2 biologically independent experiments with 4-8 mice per group. F-I are one experiment with 5-6 mice per group.

### *S. mansoni* infection modulates the myeloid transcriptome in a sex-specific manner

Since we documented significant sex dependent shifts in functional metabolism in BMDM from male and female *S. mansoni* infected mice, we sought to determine if differential transcriptional modulation underlies these shifts. In order to investigate the genes and respective pathways that were associated with specific conditions and the ones that were differentially regulated by sex we performed mRNAseq on unstimulated BMDM derived from male and female ApoE^-/-^ mice at 10-weeks post *S. mansoni* or mock infection. We identified different subsets of genes that were preferentially upregulated in males (p<0.05, Figure 6A). Among 1448 genes upregulated in males regardless of infection status with a p value <0.05, the majority of these (238 genes) were associated to metabolic functions by pathway analysis. Of these 238 genes involved in metabolism we identified hexokinase 1 (hk1), citrate synthase (cs), apolipoprotein A2 (apoa2), aldehyde dehydrogenase 3 family member A2 (aldh3a2), lipoyltransferase (lipt1), solute carrier family 19 member 1 (slc19a1), LDL receptor related protein 1 (lrp1), many of these are involved in lipoprotein and cholesterol metabolism. These data suggest that myeloid metabolism is differentially regulated by sex, at least in the context of HFD. In addition, we found 216 genes involved in immunity. Among these genes with immune function we identified interleukin 10 (il-10), Toll like receptor 5 (tlr5), NLR family pyrin domain containing 3 (nlrp3), inducible T cell costimulatory (icos), which have diverse pro and anti-inflammatory function in the immune system. Next, we surveyed the genes that are differentially regulated in males and females following *S. mansoni* infection. We found 66 genes involved in metabolism, hemostasis, the adaptive immune system, collagen degradation and not annotated to specific pathways. Following the genes with known function, the majority (10) of differentially regulated genes have documented roles in metabolism. Among these, we identified type II iodothyronine deiodinase (dio2), which has been implicated in the regulation of diet induced obesity (Kurylowicz et al., 2015; Vernia et al., 2013). Moreover, we found that hexokinase 3 (hk3), fatty acid binding protein 4 (fabp4), sphingomyelin synthase 2 (sgms2), solute carrier family 6 member 8 (slc6a8) were all upregulated in male and downregulated in female BMDM following *S. mansoni* infection (Figure 6B). In addition, we performed gene ontology analysis from the 66 genes that were differentially regulated in females and males. The most significant pathways were response to glucocorticoids, glycoprotein metabolism, and fatty acid metabolism, again suggesting that there is a sex-specific metabolic response to *S. mansoni* infection in myeloid cells. These pathways may help explain the differential metabolic modulation induced by schistosomiasis in males and females (Figure 6C).

**Figure 6.**
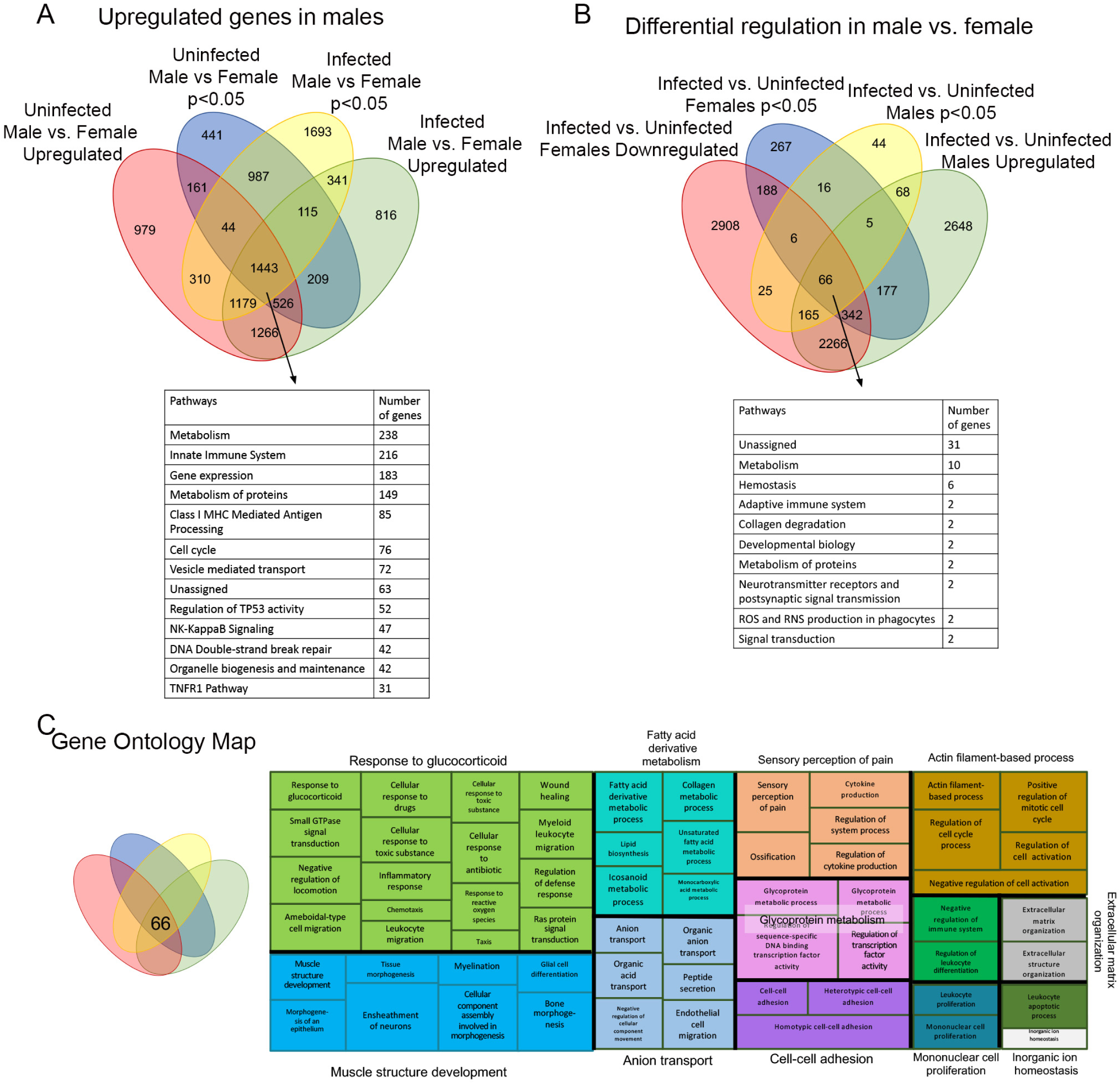
*S. mansoni* infection differentially regulates the transcriptomes of BMDM from female and male ApoE^-/-^ mice. (A) Venn diagram showing upregulated genes in males only, independently of infection state and pathway analysis corresponding to the identified genes. (B) Venn diagram and pathway analysis of differentially expressed genes in males compared to females. (C) Gene ontology map of differentially expressed genes in response to *S. mansoni* infection in male compared to female ApoE ^-/-^ mice on HFD. Sequencing data are from one experiment with 5-6 mice per group.

**Figure 7.**
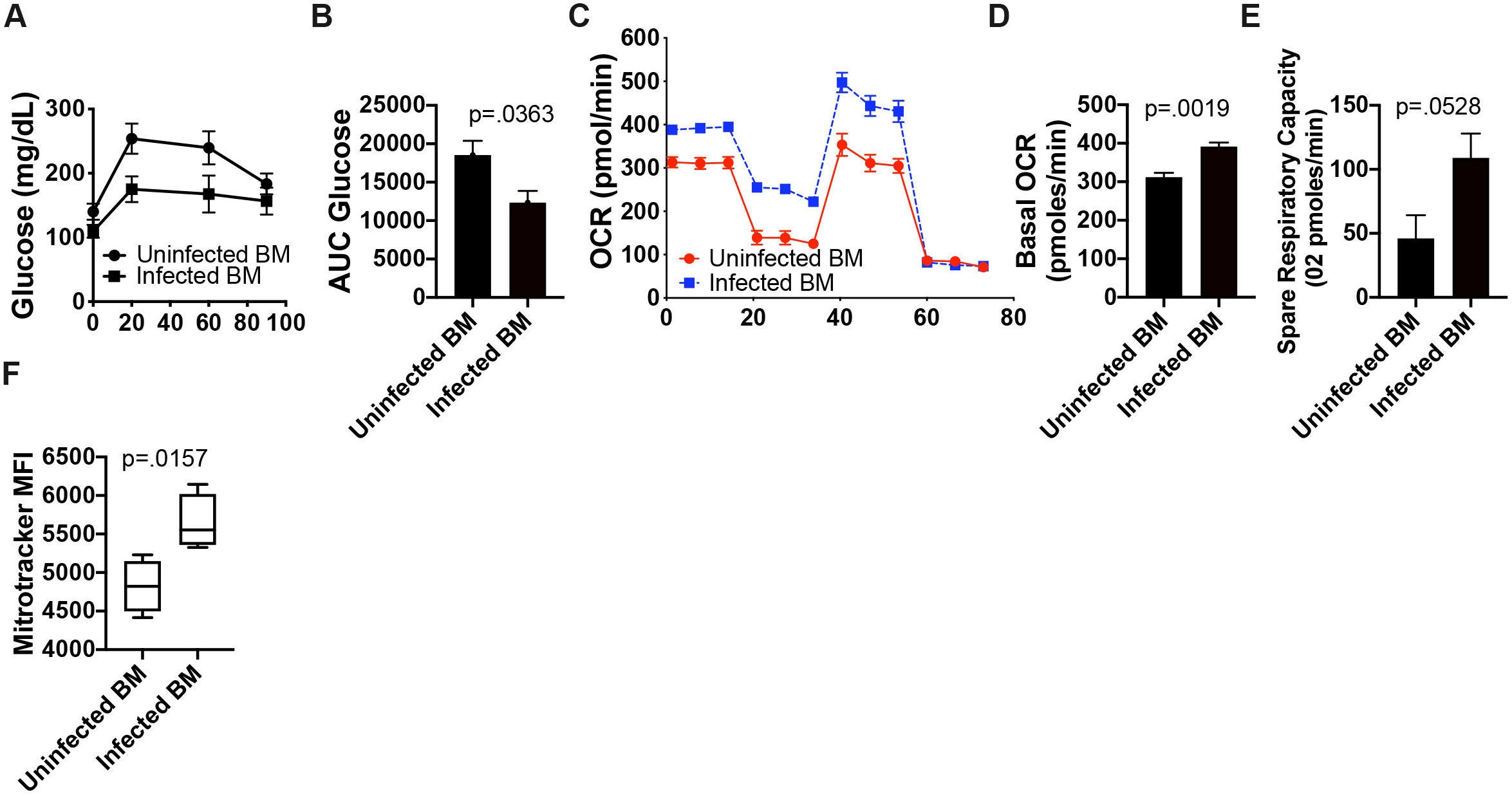
*S. mansoni* induced modulation of male macrophage metabolism is long-lived in the absence of antigen. Bone marrow from 10-week *S. mansoni* infected or control ApoE-/-mice on HFD was transferred into busulfan treated ApoE^-/-^ recipients on HFD. A,B) Glucose tolerance test (GTT) at 10 weeks post-infection. GTT values were analyzed by Area Under the Curve and graphed using GraphPad Prism. C-E) Oxygen consumption rate and spare respiratory capacity were measured at steady state in 7day BMDM. F) MitoTracker Red Deep Stain measure by flow cytometry in BMDM. Data is two combined experiments 7-8 animals per group. Statistical analysis was done using Welch’s t-tests.

### *S. mansoni* induced modulation of male macrophage metabolism is long-lived in the absence of antigen

Our metabolic and transcriptomic data from BMDM differentiated *in vitro* from male *S. mansoni* infected mice suggested that metabolic modulation may be long-lived in the absence of ongoing antigen exposure. In order to determine durable nature of modulation we transferred bone marrow from either 10-week *S. mansoni* infected male ApoE^-/-^ mice on HFD, or uninfected male controls into busulfan treated recipient ApoE^-/-^ mice on HFD. At 10 weeks post-bone marrow transfer we assayed glucose tolerance via an i.p. GTT. Mice that received bone marrow from infected males have a significantly lower glucose area under the curve (AUC) than those that received control bone marrow. We harvested bone marrow from all recipients and differentiated BMDM in M-CSF for 6 days and then performed real-time extracellular flux analysis. BMDM from recipients of bone marrow from *S. mansoni* infected mice had significantly higher basal oxygen consumption and a trend towards increased spare respiratory capacity as compared to BMDM generated from recipients of uninfected control bone marrow. Additionally, BMDM from recipients of bone marrow from *S. mansoni* infected mice had a significantly higher Mitotracker MFI than BMDM from recipients of control bone marrow. These data suggest that *S. mansoni* induced metabolic modulation of the myeloid lineage in males is long-lived even in the absence of ongoing exposure to egg antigens, and that hematopoietic cells are at least partially responsible for the regulation of whole-body glucose metabolism in the HFD ApoE^-/-^ model.

## Discussion

Helminth infections in general, and schistosomiasis in specific, have been known to be inversely correlated with obesity and glucose intolerance for over a decade, a phenomenon thought to be associated with Type 2 polarization of macrophages and T cells. In the current study we report that *S. mansoni* infection induces dramatic metabolic alterations in BMDM from male ApoE^-/-^ mice on HFD. Our results indicate that macrophages derived from the bone marrow of infected male mice have increased basal oxygen consumption and spare respiratory capacity compared to those derived from uninfected males. In T cells, an increase in spare respiratory capacity has been linked to mitochondrial biogenesis, and controls the transition to a long-lived memory phenotype (van der Windt et al., 2012). In macrophages, M2 (alternative) activation has previously been shown to lead to increased spare respiratory capacity, a process that also involves mitochondrial biogenesis (Kannan et al., 2016), while TLR recognition of bacteria has been shown to increase mitochondrial respiration via modulation of complex I and II (Garaude et al., 2016). In both cell types, the increased mitochondrial respiration underlies the longevity of the cells. In our model we have not found most of the traditional markers used to define M2 alternative activation by flow cytometry at steady state (Supplemental Figure 1). Arg1 is the only canonical M2 transcript that is modulated in the BMDM from infected male mice at steady state, and that fold increase is relatively low (.894 log FC, adjusted *p*-value =.035). Arg1 drives the production of polyamines, which in turn are able to modulate mitochondrial OxPHOS (Galvan-Pena and O’Neill, 2014; Puleston et al., 2019). In the currently accepted paradigm of M2 polarization stat6 phosphorylation upregulates PGC1-β and PPARγ, leading to mitochondrial biogenesis and increased beta-oxidation in addition to Arg1transcription. Here we present a model where neither PGC1-β or PPARγ are modulated, but Arg1 transcription is increased with a concomitant dramatic increase in mitochondrial respiration, suggesting that there may be alternative ways to modulate mitochondrial metabolism.

Alternative activation has previously been shown to be dependent on cell intrinsic lysosomal lipolysis and lal, with the defining feature being reductions in lipid droplets (Huang et al., 2014). In our model using unstimulated BMDM from *S. mansoni* infected mice there is no gross difference in lipid droplets, nor an increase in lal transcripts. Instead, we found a significant shift in the lipidome of BMDM from infected male mice that centered on a reduction in cholesterol esters. Previously published work in IL-4 induced M2 macrophages has found that lipolysis centered on TGs as a fatty acid source. While we did find reductions in two species of DGs and one species of monoacylglycerol (MG), there were no significant alterations to TAGs in the BMDM generated from infected males. Flux analysis with heavy carbon labeled glucose and palmitate suggest that BMDM from male infected mice have increased shuttling of both glucose and palmitate into TCA cycle intermediates, suggesting that these cells have increased mitochondrial beta oxidation. In addition to the unique lipidomic modulations, we found that the dramatic increase in both basal oxygen consumption and spare respiratory capacity we observed in BMDM from male infected mice was significantly palmitate dependent. This observation further supports the idea that *in vivo* exposure of myeloid precursors to helminth antigens induces unique metabolic modulations that focus on cholesterol and lipid metabolism as a source for palmitate for beta oxidation. These data suggest that hybrid metabolic states in the absence of M1 or M2 polarization occur in macrophages in the context of chronic disease, and present a challenge to the dichotomy of M1 versus M2 activation being driven by immunometabolism. Future studies exploring the immunometabolism of bone marrow derived and tissue resident macrophages and dendritic cells from other chronic infections and inflammatory diseases are needed in order to obtain a true picture of the correlation between metabolism and myeloid polarization.

There are significant clinical differences in both the etiology and pathology of diabetes and cardiovascular disease between males and females, but sex differences in immunological activation or modulation by *S. mansoni* infection have not previously been studied in humans or animal models. Surprisingly, we found that the unique metabolic modulations induced by *S. mansoni* infection in BMDM from male mice do not occur in females, and infected females are not reliably protected from high-fat diet induced obesity or glucose intolerance. Interestingly, hepatic macrophage alternative activation in response to infection is equivalent between males and females, but infection increases blood monocyte frequency in females to a much greater extent than in males, suggesting sex specific modulation of either monopoiesis or monocyte recruitment into tissues. Blood monocytes from infected males phenocopy the increase in Mitotracker MFI that we have found in BMDM generated from infected males, indicating that our BMDM model is likely an accurate representation of the *in vivo* potential of the myeloid compartment. Interestingly, we found that infection significantly reduced the total number of CMP and GMP in male, but not female bone marrow, again pointing to a sex-specific modulation of the myeloid lineage at the precursor level. The reduction of CMP and GMP in infected males suggests either an increase in the rate of differentiation into monocytes/granulocytes, or decreased homeostatic proliferation. Since males have fewer blood monocytes than female, we favor the later possibility. These possibilities and the relationship between reductions in numbers of CMP and GMP in males, and the functional metabolic differences in BMDM will be explored in future work.

Analysis of the BMDM transcriptomes from males and females revealed that over a thousand genes are up regulated in male versus female BMDM regardless of *S. mansoni* infection state. At the same time, over five dozen are differentially regulated by infection. Focusing on the genes that are up-regulated in BMDM from infected males and down-regulated in females, we found more than ten genes with known functions in cellular metabolism. Some of these genes, like PFKFB3, have known regulatory elements for progesterone and estrogen (Shi et al., 2017), but some of them, like fabp4, have no published mechanism of sex hormone regulation. Glycolysis and cholesterol metabolism have previously been shown to directly affect the inflammatory potential of myeloid cells. Our data indicate that *S. mansoni* infection induces a hybrid inflammatory state in male but not female BMDM, where the LPS induced production of nitrite and iNOS is enhanced, while the production of the key chronic pro-inflammatory mediators IL-12p70, CXCL1, and IL-6 are reduced. This inflammatory profile is distinct from what has previously been published with IL-4 induced M2 macrophages, where the production of iNOS and nitrite are reduced following TLR stimulation (Lam et al., 2016). CXCL1 and IL-6 have previously been linked to increased monocyte recruitment and disease progression in both atherosclerosis and obesity-induced diabetes (Boisvert et al., 2006; Hartman and Frishman, 2014; Nunemaker et al., 2014; Qu et al., 2014; Zhou et al., 2011), so these data support the possibility of infection driven modulation of macrophage function supporting the decreased pathology seen in infected males. We have demonstrated that the modulation of macrophage oxygen consumption is transferrable to an uninfected recipient, and can last for at least ten weeks, suggesting that in males, *S. mansoni* infection induces long-lived metabolic modulation of the myeloid lineage that survives in the absence of ongoing antigenic exposure. Trained innate immunity has previously been documented to be induced by BCG immunization (Kaufmann et al., 2018), and has recently been suggested to be the mechanism underlying the association between previous bacterial and fungal infections and the development of atherosclerosis (Hoogeveen et al., 2018; Leentjens et al., 2018). In these models, trained immunity and epigenetic reprogramming is driven in part from a switch from oxidative phosphorylation to increased aerobic glycolysis (Cheng et al., 2014; Stienstra et al., 2017). In the case of BCG, trained circulating monocytes can be found months after immunization, which strongly suggests reprogramming of bone marrow progenitors (Ifrim et al., 2014). Recent reports indicate that western high-fat diet itself also induces innate training of bone marrow progenitors in both the Ldr^-/-^ model of atherosclerosis (Christ et al., 2018) and in obesity related steatohepatitis (Krenkel et al., 2020). Our data suggests that *S. mansoni* infection in males trains the myeloid lineage in the opposite fashion; modulating the metabolic transcriptome of the myeloid lineage such that oxidative phosphorylation and mitochondrial activity is increased, while the chronic inflammatory potential is decreased. Alterations to the numbers and frequency of myeloid progenitors and the transferability of our phenotype via bone marrow suggests that progenitors are indeed modulated in our model, the relative role of epigenetic modulation versus microRNA regulation in dictating the *S. mansoni* induced myeloid transcriptome will be the subject of future studies.

There are significant differences in both the susceptibility and disease presentation between males and females for both diabetes and cardiovascular disease, but few studies have focused on the role of immunometabolism in this dichotomy. Our data suggests apparent biological sex-dependent differences in both the ability of schistosomes to protect from the development of HFD induced metabolic disease parameters, and the ability to modulate macrophage glucose and lipid metabolism. The current epidemiological data strongly indicates an inverse correlation between helminth infections and metabolic diseases such as diabetes and cardiovascular disease, but these studies were not designed to identify sex-specific correlations. Few animal studies have focused on the role of myeloid cells in driving sex-specific metabolic differences. Our current data indicate a clear need for further studies in both humans and animal models to specifically probe the relationship between biological sex and myeloid metabolism, and how chronic helminth infections modulate this.

## Acknowledgements

The work was supported by The University of Utah, a Scientist Development Grant from the American Heart Association to KCF (14SDG18230012), an American Heart Association Pre-doctoral Award (18PRE34030086) to DCS, and 1R21AI135385-01A1 to EA. *B. glabrata* snails provided by the NIAID Schistosomiasis Resource Center of the Biomedical Research Institute (Rockville, MD) through NIH-NIAID Contract HHSN272201700014I for distribution through BEI Resources. JEC is supported through U54 DK110858-01, mass spectrometry equipment employed was provided by 1S10OD016232-01, 1S10OD018210-01A1 and 1S10OD021505-01 to JEC.

## STAR Methods

### KEY RESOURCES TABLE

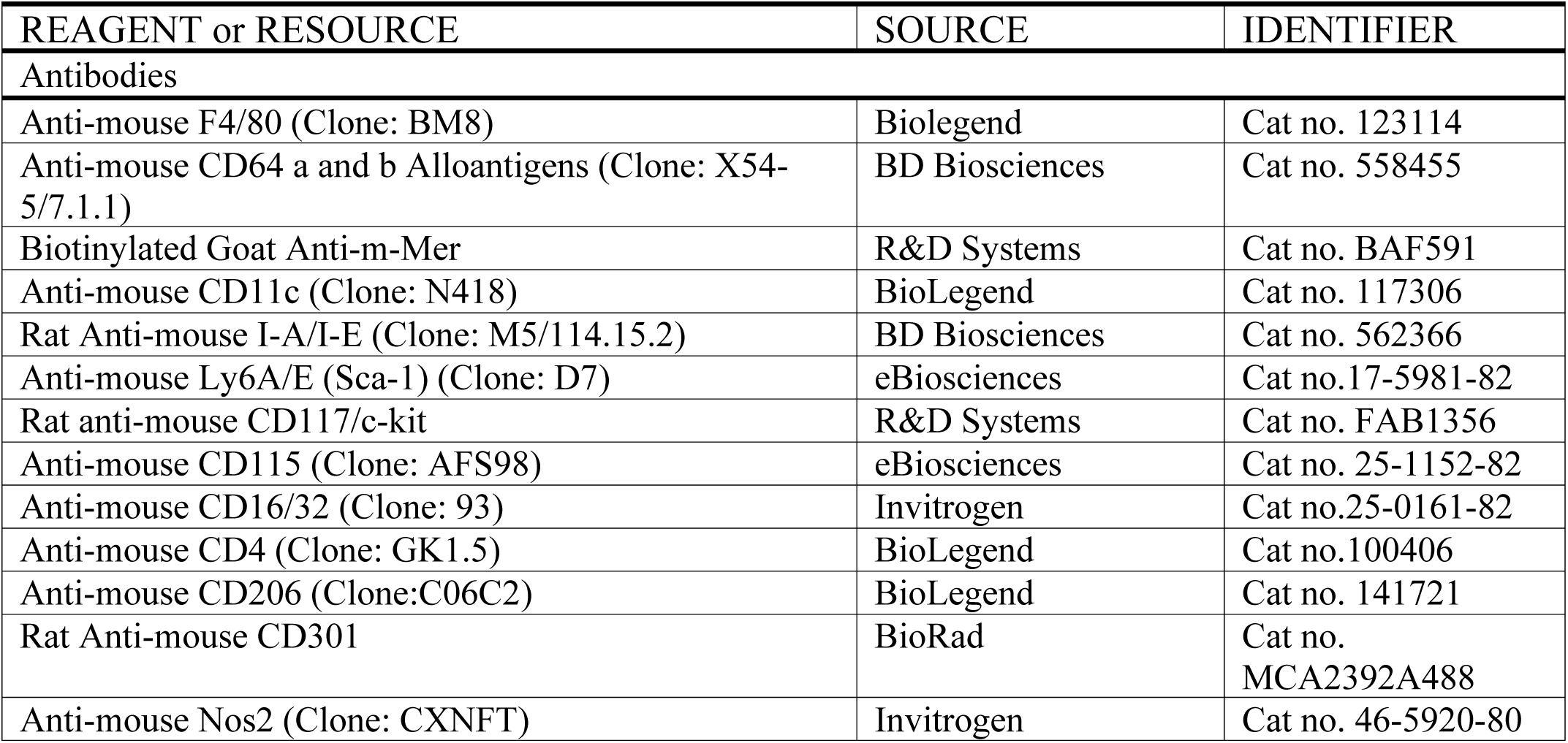
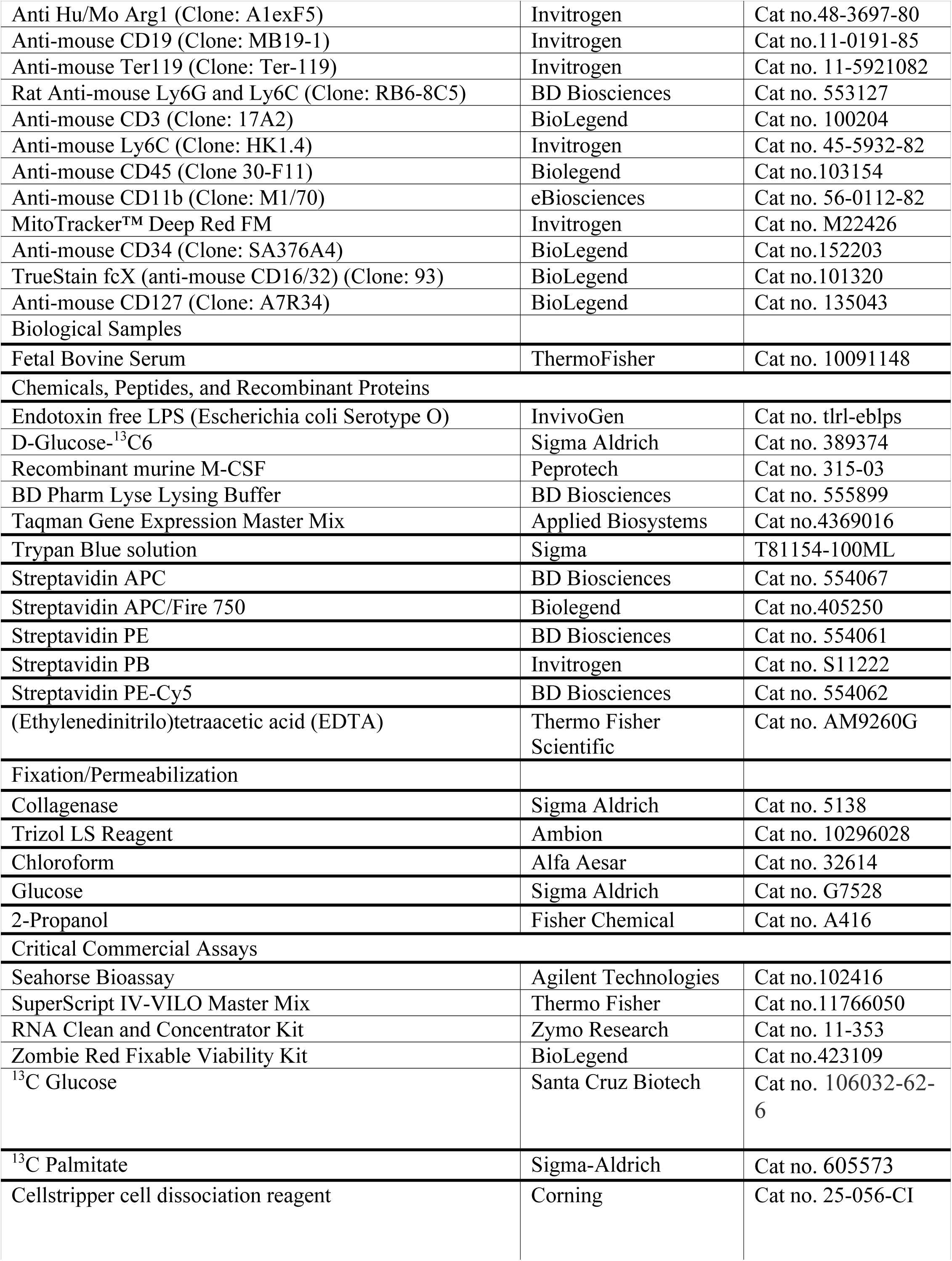
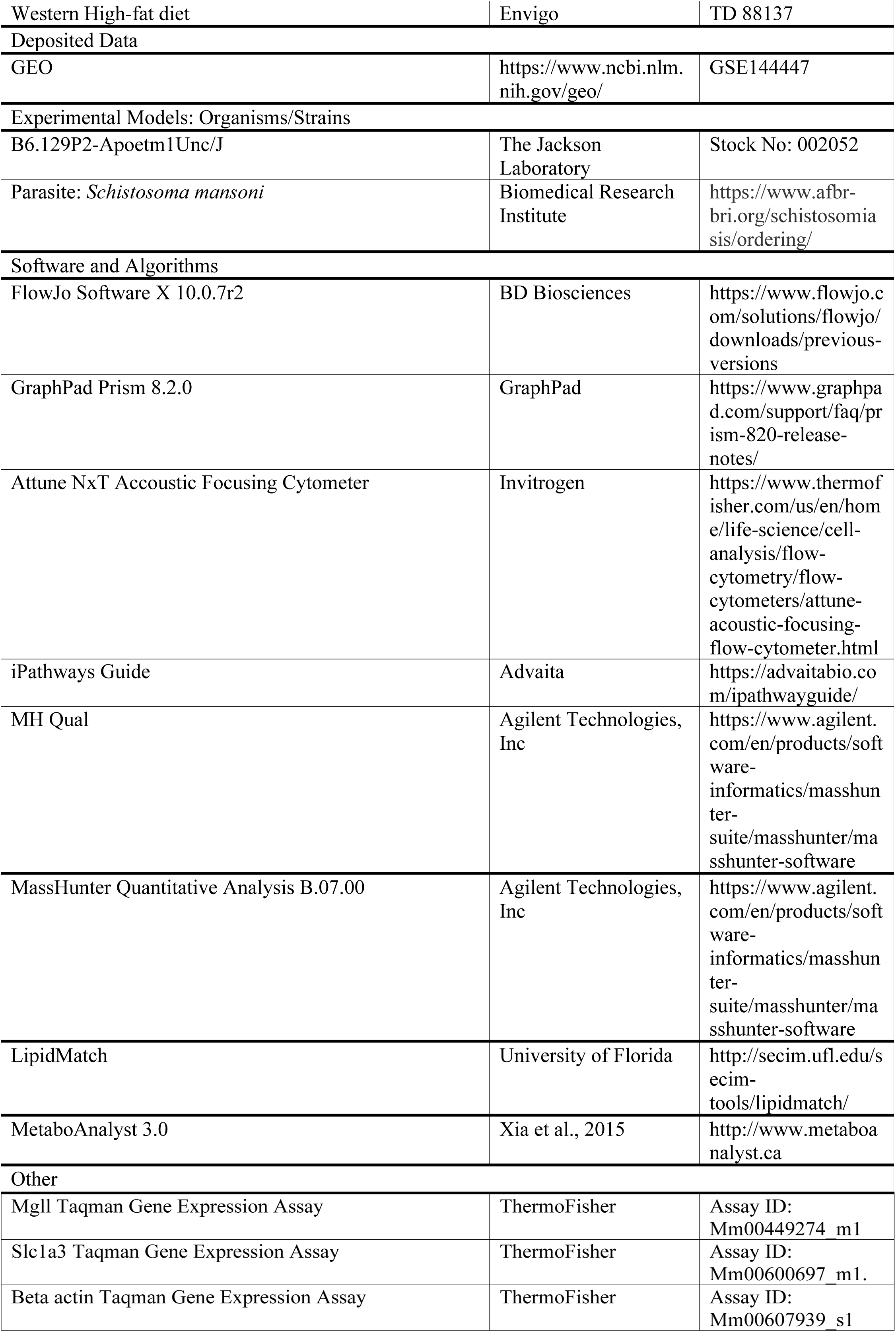

### LEAD CONTACT AND MATERIALS AVAILABILITY

Further information requests should be directed to and will be fulfilled by the Lead Contact, Keke Fairfax. These studies generated no new reagents.

### EXPERIMENTAL MODEL AND SUBJECT DETAILS

#### Parasite and Mouse Models

This study was carried out in accordance with the recommendations in the Guide for the Care and Use of Laboratory Animals of the National Institutes of Health. The protocols were approved by the Institutional Animal Care and Use Committees of the University of Utah and Purdue University. Snails infected with *S. mansoni* (strain NMRI, NR-21962) were provided by the Schistosome Research Reagent Resource Center for distribution by BEI Resources, NIAID NIH. Male and female ApoE^-/-^ (B6.129P2-Apoetm1Unc/J) were purchased from the Jackson Laboratories and bred at the University of Utah. Mice were housed in pathogen-free conditions and were fed standard rodent chow (2019 rodent chow, Harlan Teklad) until 10-14 days before infection when they were transitioned to a high-fat diet (HFD: 21% milk fat, 0.15% cholesterol: TD 88137 Envigo). Bone marrow chimeras were generated by treating ApoE^-/-^ mice that had been on high-fat diet for 4 weeks with 20mg/kg of pharmaceutical grade busulfan for 5 days (total dose of 100mg/kg). On day 6 mice were i.v. injected with 2.5-3 × 10^6^ bone marrow cells from either 10-week *S. mansoni* infected or control uninfected ApoE^-/-^ mice on high-fat diet. Reconstitution was validated via flow-cytometry at 3-weeks post-transfer and recipient mice were maintained on high-fat diet for 10-weeks post-reconstitution.

### METHOD DETAILS

#### *S. mansoni* infection and glucose tolerance test

ApoE^-/-^ female and male mice of 6 weeks of age were exposed percutaneously to 75-90 cercariae of *S. mansoni* or were mocked infected (as controls). At five- and ten-weeks post-infection mice were fasted for 5 hours and baseline blood glucose levels were obtained via lateral tail vein nick. Mice were then administered a single intraperitoneal injection of glucose (2mg/g of body weight, ultrapure glucose, Sigma G7528). Blood glucose levels were obtained at 20, 60, and 90 min post injection. Individual data points obtained were analyzed by Area Under Curve (AUC).

#### Mouse macrophage culture

Mouse bone marrow-derived macrophages (BMDM) were generated as follows: bone marrow cells were isolated by centrifugation of bones at >10,000 x g in a microcentrifuge tube for 15 seconds as previously described (Amend et al., 2016). Cells were differentiated in M-CSF (20ng/mL, Peprotech, Rocky Hill, NJ) in complete macrophage medium (CMM: RPMI1640, 10% FCS, 2mM L-glutamine and 1 IU/mL Pen-Strep for 6 or 7 days. On the last day, cells were harvested in Cellstripper cell dissociation reagent (Corning) were washed with CMM) and prepared for downstream assays.

#### Glycolytic and phospho-oxidative metabolism measurement (seahorse assay)

BMDM from different conditions (uninfected controls or *S. mansoni* infected) were resuspended at the same concentration in XF assay media supplemented with 5% FCS and 5mM glucose. The day before the assay, the probe plate was calibrated and incubated at 37 C in a non-CO2 incubator. Resuspended cells were seeded at a concentration of 1.5 × 10^5^ cells per well and incubated for 20-60 minutes in the Prep Station incubator (37 C non-CO2 incubator). Following initial incubation, XF Running Media (XF assay media with 5% FCS and 10mM Glucose) were dispensed into each well. OCR and ECAR were measured by an XF96 Seahorse Extracellular Flux Analyzer following the manufacturer’s instructions. For the seahorse assay, cells were treated with oligomycin (1uM), FCCP (1.5uM), rotenone (100nM) and antimycin A (1uM). Each condition was performed in 2-3 technical replicates. For determination of palmitate dependent respiration, BSA-conjugated palmitate (BSA: palmitate = 1:6, molar ratio) was prepared according to the Seahorse protocol (Seahorse Bioscience). Briefly, 1 mM sodium palmitate (Sigma Aldrich) was conjugated with 0.17 mM fatty acid free-BSA (Sigma Aldrich) in 150 mM NaCl solution at 37°C for 1h. Palmitate-BSA was stored in glass vials at −20°C until use. Cells were incubated as above in glucose limited XF media per manufacturer instructions.

#### Flow Cytometry

Livers from uninfected and infected mice were perfused with 1X PBS, mashed and digested in DMEM 0.5% Collagenase (Sigma) at 37°C for 15 min. Then, livers were mashed and filtered through a 100 μm metal strainer and digested an additional 15 min. Following second digestion, the liver contents were strained, washed with DMEM and spun down 1500 rpm for 5 min. The pellet was lysed with 1X lysis buffer (BD PharmLyse), quenched with 1% FBS DMEM, and washed to be used in flow cytometry. Surface staining was performed using the following mAb against mouse antigens: CD45 (30-F11, eBioscience), CD301(BioRad), CD206 (C068C2, Biolegend), F4/80 (BM8, Biolegend), mouse Mer biotinylated (R&D), CD64(X54-5/7.1, BD). Intracellular antigen staining such as Nos2 (CXNFT, Invitrogen, and C-11, Santa Cruz) and Arg1 (Invitrogen, A1exF5)) was performed using the Intracellular Fixation and Permeabilization Buffer set (Thermo Fisher Scientific cat. no. 88-8824) per manufacturer’s instruction. Further, bone marrow cells were obtained by centrifugation of bones into tubes at >10000 rpm for 15 s. Surface staining for bone marrow precursors was performed using the following antibodies: Ter119, CD19, CD4, CD3, Gr-1, CD11b, Sca-1, CD115, Ly6C, c-Kit, flt3, CD127 and CD16/32. PBMC from whole blood were obtained following red blood cell lysis and used for flow cytometry analysis. Surface staining of PBMC was performed using Ter119, CD64, CD11b, CD115, Ly6C and MitoTracker Red.

Samples were acquired using Attune NxT Focusing Flow Cytometer (Thermo Fisher Scientific) and analyzed using Flowjo X 10.0.7r2 (FlowJo LLC, Inc.).

#### RNA Isolation and q-RT-PCR

BMDM were stored in Trizol, and RNA isolation was performed as described in the Immunological Genome Project Total RNA isolation protocol. Next, cDNA was synthesized from RNA using Superscript IV VILO (ThermoFisher Scientific) for reverse transcription. qPCR was performed using TaqMan Gene expression assays (Mgll, Slc1a3, beta actin, ThermoFisher) on an Applied Biosystems Stepone Plus Real-Time PCR System. Beta-Actin assay number Mm00607939_s1, mgll assay Mm00449274_m1, slc1a3 assay Mm00600697_m1. Relative expression was calculated using the 2-ΔΔCt method.

#### Measurement of Cytokines and Inflammatory mediators

For cytokine levels of BMDCs, supernatants were collected at 24 hours post stimulation and measured with Mouse Cytokine and Chemokine ProcartaPlex 26plex panel (Life Technologies) per manufacture instructions using a Luminex Magpix system. Nitrite levels in cell culture media were determined using a Griess reagent kit for nitrite determination (Invitrogen) according the manufacturer’s instructions.

#### Untargeted lipidomics

##### Sample extraction from serum or cell pellets

Lipids were extracted from serum (50μL) or cell pellets in a combined solution as described in (Matyash et al., 2008). In detail, samples were combined in solution with 225 μL MeOH containing internal standards (IS; Avanti splash Lipidomix (Lot#12), 10μL each sample) and 750 μL methyl tert-butyl ether (MTBE). The samples were sonicated for 1 min, rested on ice for 1 hour, briefly vortexed every 15 min then an addition of 200μL dd-H2O was made to induce phase separation. All solutions were pre-chilled on ice. The sample were then vortexed for 20 s, rested at room temperature for 10 min, and centrifuged at 14,000 g for 10 min at 4 C. The upper organic phase was collected and evaporated to dryness under vacuum. Lipid samples were reconstituted in 200μL IPA and transferred to an LC-MS vial with insert for analysis.

Concurrently, a process blank sample was brought forward as well as quality control sample was prepared by taking equal volumes (10μL per sample) from each sample after final resuspension.

##### LC-MS Methods

Lipid extracts were separated on a Waters Acquity UPLC CSH C18 1.7 um 2.1 × 100 mm column maintained at 65 °C connected to an Agilent HiP 1290 Sampler, Agilent 1290 Infinity pump, equipped with an Agilent 1290 Flex Cube and Agilent 6530 Accurate Mass Q-TOF dual ESI mass spectrometer. For positive mode, the source gas temperature was set to 225 °C, with a gas flow of 11 L/min and a nebulizer pressure of 50 psig. VCap voltage was set at 3500 V, fragmentor at 110 V, skimmer at 85 V and Octopole RF peak at 750 V. For negative mode, the source gas temperature was set at 325 °C, with a drying gas flow of 12 L/min and a nebulizer pressure of 30 psig. VCap voltage is set 3000 V, fragmentor at125 V, skimmer at 75 V and Octopole RF peak at 750 V. Reference masses in positive mode (*m/z* 121.0509 and 922.0098) were infused with nebulizer pressure at 2 psig, in negative mode reference masses (*m/z* 1033.988, 966.0007, 112.9856 and 68.9958) were infused with a nebulizer pressure at 5psig. Samples were analyzed in a randomized order in both positive and negative ionization modes in separate experiments acquiring with the scan range m/z 100-1700. Mobile phase A consisted of ACN:H_2_O (60:40 *v/v*) in 10 mM ammonium formate and 0.1% formic acid, and mobile phase B consisted of IPA:ACN:H_2_0 (90:9:1 *v/v*) in 10 mM ammonium formate and 0.1% formic acid. The chromatography gradient for both positive and negative modes started at 15% mobile phase B then increased to 30% B over 2.4 min, then increased to 48% from 2.4-3.0 min, followed by an increase to 82% B from 3-13.2 min, and then to 99% from 13.2-13.8 min where it was held until 15.4 min and then returned to the initial conditioned and equilibrated for 4 min. Flow was 0.5 mL/min throughout, injection volume was 5μL for positive and 7 μL negative mode. Tandem mass spectrometry is conducted using the same LC gradient at collision energies of 20 V and 40 V.

#### Targeted lipidomics

##### Sample Preparation

Lipids were extracted from cell pellets (500,000 cells) as described in detail above (Matyash et al., 2008). For targeted lipidomics, lipid extracts were separated on a Waters BEH HILIC column 1.7 μm, 100 mm × 3 mm column maintained at 60 °C connected to an Agilent HiP 1290 Sampler, Agilent 1290 Infinity pump, and equipped with an Agilent 6490 triple quadrupole (QqQ) mass spectrometer. Lipids were detected using dynamic multiple reaction monitoring (dMRM) in negative ion mode. Source gas temperature is set to 225 °C, with a gas flow of 13 L/min and a nebulizer pressure of 30 psi. Sheath gas temperature was 350 °C, sheath gas flow was 11 L/min, capillary voltage of 4000 V, nozzle voltage was 500 V, high pressure RF was 190 V and low-pressure RF was 120 V. Injection volume was 2 µL and the samples were injected in a randomized order. Mobile phase A consisted of H_2_O, mobile phase B consisted of ACN:H_2_O (96:4 *v/v*) and both contain 7 mM ammonium acetate. The chromatography gradient started at 100% mobile phase B and decreased to 84% B over 10 min. Post-time was 9 min and the flow rate was 0.4 mL/min throughout. Collision energies (25 V) and cell accelerator voltages (3 V) were optimized using lipid standards with dMRM quantifier transitions as [M-H]^-^→[T2_RCOO]^-^ and qualifier transitions of [M-H]^-^→[T1_RCOO]^-^ and [M-H]^-^→[NL_T2Ketene]^-^.

#### Lipid data analysis

The pooled QC samples and process blank samples were injected throughout the sample queue to ensure the reliability of acquired LC-MS data. Results from LC-MS experiments were collected using Agilent MassHunter (MH) Workstation and analyzed using the software packages MH Qual, MH Quant (Agilent Technologies, Inc) and LipidMatch to prepare the data set. The data table exported from MHQuant was evaluated using Excel where initial lipid targets were parsed based on the following criteria. Only lipids with relative standard deviation (RSD) less than 30% in QC samples were used for data analysis. Additionally, targets identified in blanks or double blanks at significant amounts (area under the curve (AUC) target blank/AUC target QC >30%) were removed from analysis. Lipids were quantitated based on peak area ratios to the spiked IS of the same or nearest class.

#### RNA Sequencing

##### Library Preparation and sequencing

The concentration and quality of total RNA samples was first assessed using Agilent 2100 Bioanalyzer. A RIN (RNA Integrity Number) of five or higher was required to pass the quality control. Then 200 nanograms of RNA per sample were used to prepared dual-indexed strand-specific cDNA library using KAPA mRNA Hyperprep Kit (Roche). The resulting libraries were assessed for its quantity and size distribution using Qubit and Agilent 2100 Bioanalyzer. Two hundred pico molar pooled libraries were utilized per flowcell for clustering amplification on cBot using HiSeq 3000/4000 PE Cluster Kit and sequenced with 2.75bp paired-end configuration on HiSeq4000 (Illumina) using HiSeq 3000/4000 PE SBS Kit. A Phred quality score (Q score) was used to measure the quality of sequencing. More than 90% of the sequencing reads reached Q30 (99.9% base call accuracy).

##### Sequence alignment and gene counts

The sequencing data were first assessed using FastQC (Babraham Bioinformatics, Cambridge, UK) for quality control. Then all sequenced libraries were mapped to the mouse genome (UCSC mm10) using STAR RNA-seq aligner (Dobin et al., 2013) with the following parameter: “-- outSAMmapqUnique 60”. The reads distribution across the genome was assessed using bamutils (from ngsutils) (Breese and Liu, 2013). Uniquely mapped sequencing reads were assigned to mm10 refGene genes using featureCounts (from subread) (Liao et al., 2014) with the following parameters: “-s 2 –p –Q 10”. Quality control of sequencing and mapping results was summarized using MultiQC (Ewels et al., 2016). Genes with read count per million (CPM) > 0.5 in more than 2 of the samples were kept. The data was normalized using TMM (trimmed mean of M values) method. Differential expression analysis was performed using edgeR (McCarthy et al., 2012; Robinson et al., 2010). False discovery rate (FDR) was computed from p-values using the Benjamini-Hochberg procedure.

##### Pathway analysis

The Data (significantly impacted pathways, biological processes, molecular interactions.) were analyzed using Advaita Bio’s iPathwayGuide (http://www.advaitabio.com/ipathwayguide). Pathway analysis was performed on log_2_-transformed data using Bonferroni-corrected *p*-values. The data discussed in this publication have been deposited in NCBI’s Gene Expression Omnibus (Edgar *et al*., 2002) and are accessible through GEO Series accession number GSE 144447(https://www.ncbi.nlm.nih.gov/geo/query/acc.cgi?acc=GSE144447)

#### Heavy glucose and heavy palmitate labeling

For metabolomics tracing in Figure 2 BMDM were differentiated in CMM containing normal glucose. At Day 6 of culture cells were switched to CMM containing ^13^C_6_-glucose (Santa Cruz Biotech) for 24 hours. Cells were harvested and processed as described below. For metabolomics tracing in figure 5 BMDM were differentiated in CMM containing normal glucose and serum. At Day 6 of culture cells were switched to CMM containing dialyzed serum and 1 mM ^13^C palmitate (sigma Aldrich) for 36 hours. Cells were harvested and processed as described below.

#### Metabolomics

##### Extraction

Cold 90% methanol (MeOH) solution was added to each sample to give a final concentration of 80% MeOH to each cell pellet. Samples were incubated at −20 °C for 1 hr. After incubation, the samples were centrifuged at 20,000 x g for 10 minutes at 4 °C. The supernatant was transferred from each sample tube into a labeled, fresh micro centrifuge tube. The samples were dried *en vacuo*.

##### Mass Spectrometry Analysis of Samples

All GC-MS analysis was performed with an Agilent 7200 GC-QTOF and an Agilent 7693A automatic liquid sampler. Dried samples were suspended in 40 µL of a 40 mg/mL O-methoxylamine hydrochloride (MOX) (MP Bio #155405) in dry pyridine (EMD Millipore #PX2012-7) and incubated for one hour at 37 °C in a sand bath. 25 µL of this solution was added to auto sampler vials followed by the automatic addition of 60 µL of N-methyl-N-trimethylsilyltrifluoracetamide (MSTFA with 1%TMCS, Thermo #TS48913) and incubated for 30 minutes at 37 °C. Following incubation, each sample were vortexed and 1 µL of the prepared sample was injected into the gas chromatograph inlet in the split mode with the inlet temperature held at 250 °C. A 10:1 split ratio was used for analysis for the majority of metabolites. Any metabolites that saturated the instrument at the 10:1 split was analyzed at a 50:1 split ratio. The gas chromatograph had an initial temperature of 60 °C for one minute followed by a 10°C/min ramp to 325 °C and a hold time of 10 minutes. A 30-meter Agilent Zorbax DB-5MS with 10 m Duraguard capillary column was employed for chromatographic separation. Helium was used as the carrier gas at a rate of 1 mL/min.

##### Data Analysis

The area under the curve for each isotope was extracted using MHQuant software (Agilent). This data was exported as a .csv file and isotopically corrected using an in house modified version of DeuteRater (Naylor et al., 2017)

#### Machine Learning

The selection of the most informative or important features (i.e., the features contributing to the prediction) was performed using a machine-learning approach involving an elastic-net regressor, which followed a round of traditional univariate filtering. The process included two steps:

1. Creating a series of ANOVA models (one for each of the lipid-based features), and pre-selecting features based on η^2^ to limit the complexity of the downstream elastic net model. The uninformative features were rejected and not used in the second step.
2. Establishing an elastic net regression model via cross-validation and grid-search of the parameters. The zero coefficients of the model were removed. The top 20 non-zero coefficients provided ranking for the features in terms of their importance.

The elastic net regression attempts to minimize the following functional:

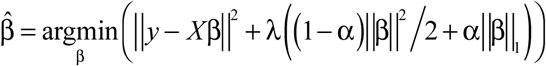

over a grid of α and λ values (Zou, 2005). The elastic-net approach linearly combines the L_1_ and L_2_ penalties used in LASSO (Tibshirani, 1996) and ridge-regression methods (Hoerl, 1970), respectively. Therefore, the elastic net penalty would become LASSO penalty for α = 1 and ridge penalty for α = 0. The parameter λ controls the overall strength of the combined penalty term. The input *X* in the model consists of all the molecular features identified, and y is probability of observing a particular animal.

#### Statistical Analysis

Statistical analyses of data were performed using one-way ANOVA, a non-parametric Mann-Whitney test, or unpaired Student’s t-test depending on the data distribution. P ≤ 0.05 were considered statistically significant. Analyses and graphing were performed using Prism (GraphPad v8.0) and R-language for statistical computing.

