## Supplemental figures for "*Schistosoma mansoni* infection reprograms the metabolic potential of the myeloid lineage in a mouse model of metabolic syndrome"

### Supplementary Figure 1

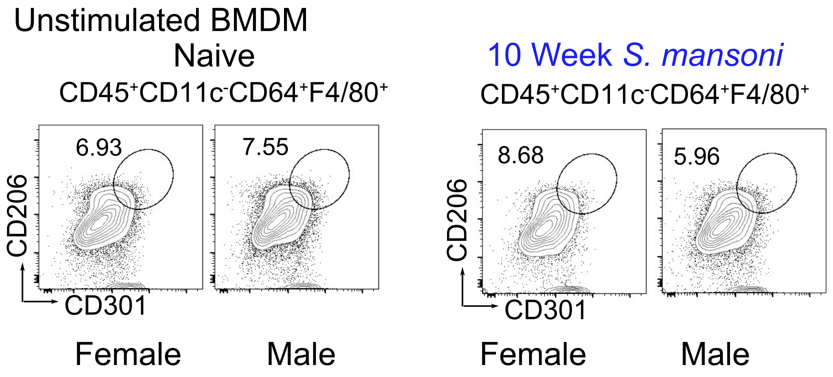


**Supplementary Figure 1. BMDM derived from *S. mansoni* infected males and females are not alternatively activated.** (A) Unstimulated BMDM from naive ApoE-/- mice on HFD. (B) Unstimulated BMDM from 10 week S. mansoni infected ApoE-/- mice.

A

| Molecular feature | Coefficient | Importance score | Effect size (eta squared) | | | | | | | Adjusted p-value | | | | | | |
|---|---|---|---|---|---|---|---|---|---|---|---|---|---|---|---|---|
| | | | Infection | Sex | Media | Infection:Sex | Infection:Media | Sex:Media | Infection:Sex:Media | Infection | Sex | Media | Infection:Sex | Infection:Media | Sex:Media | Infection:Sex:Media |
| PG 18:1_18:2 | 0.4428 | 100.0000 | 0.0288 | 0.2030 | 0.0261 | 0.1177 | 0.0233 | 0.0449 | 0.0000 | 0.1323 | 0.8898 | 0.7250 | 0.3622 | 0.8161 | 0.9745 | 0.9972 |
| PG 18:1_22:5 | 0.2995 | 67.6219 | 0.0508 | 0.1274 | 0.0298 | 0.2653 | 0.0444 | 0.0009 | 0.0057 | 0.0171 | 0.7500 | 0.3728 | 0.0972 | 0.5891 | 0.9745 | 0.9967 |
| PG 16:0_18:1 | 0.2943 | 66.4469 | 0.0344 | 0.2386 | 0.3957 | 0.2535 | 0.0458 | 0.0051 | 0.0389 | 0.0112 | 0.8654 | 0.0343 | 0.0513 | 0.4217 | 0.9745 | 0.9537 |
| BMP 16:0_18:1 | -0.2884 | 65.1314 | 0.1651 | 0.0946 | 0.5496 | 0.4106 | 0.0740 | 0.0754 | 0.0766 | 0.0010 | 0.0043 | 0.0005 | 0.0152 | 0.4217 | 0.7370 | 0.9537 |
| PG 20:4_20:4 | -0.2446 | 55.2417 | 0.0305 | 0.0290 | 0.0004 | 0.3707 | 0.0160 | 0.0659 | 0.0291 | 0.0484 | 0.0790 | 0.7815 | 0.1205 | 0.9062 | 0.9745 | 0.9537 |
| PG 16:0_18:2 | 0.2203 | 49.7407 | 0.0005 | 0.2169 | 0.7788 | 0.2178 | 0.0156 | 0.0212 | 0.0000 | 0.1303 | 0.8654 | 0.0005 | 0.1646 | 0.8567 | 0.9745 | 0.9972 |
| BMP 22:4_22:5 | 0.2186 | 49.3521 | 0.0034 | 0.1335 | 0.0454 | 0.0042 | 0.0043 | 0.0680 | 0.0164 | 0.7241 | 0.9117 | 0.6433 | 0.8835 | 0.7354 | 0.9142 | 0.9967 |
| LPE 18:1 d7 | 0.1650 | 37.2621 | 0.0484 | 0.1412 | 0.2852 | 0.0053 | 0.0154 | 0.0208 | 0.0013 | 0.8895 | 0.8654 | 0.1585 | 0.9068 | 0.9062 | 0.9745 | 0.9972 |
| PI 15:0_18:1 d7 | -0.1315 | 29.6880 | 0.0502 | 0.0271 | 0.0046 | 0.0001 | 0.0055 | 0.0003 | 0.0031 | 0.8895 | 0.8654 | 0.7815 | 0.9013 | 0.8564 | 0.9745 | 0.9967 |
| PG 20:3_22:6 | -0.1152 | 26.0116 | 0.2722 | 0.0291 | 0.1021 | 0.0363 | 0.0338 | 0.0132 | 0.0091 | 0.0262 | 0.5341 | 0.1871 | 0.4458 | 0.5891 | 0.9745 | 0.9967 |
| BMP 22:6_22:6 | 0.0972 | 21.9517 | 0.2267 | 0.1593 | 0.2785 | 0.0962 | 0.0036 | 0.0063 | 0.0738 | 0.0112 | 0.8319 | 0.9555 | 0.1205 | 0.4948 | 0.9142 | 0.9537 |
| BMP 18:1_22:4 | -0.0882 | 19.9204 | 0.1333 | 0.0060 | 0.2395 | 0.3092 | 0.0221 | 0.0895 | 0.0546 | 0.0034 | 0.0301 | 0.0159 | 0.0235 | 0.4409 | 0.7370 | 0.9537 |
| PG 16:1_18:2 | 0.0844 | 19.0487 | 0.0196 | 0.1398 | 0.1288 | 0.0447 | 0.0820 | 0.0154 | 0.0081 | 0.5644 | 0.8919 | 0.9555 | 0.7790 | 0.7646 | 0.9745 | 0.9967 |
| PG 20:3_20:4 | -0.0762 | 17.2121 | 0.0032 | 0.0351 | 0.5305 | 0.0971 | 0.0043 | 0.1257 | 0.0057 | 0.4312 | 0.2361 | 0.0154 | 0.4879 | 0.9873 | 0.9213 | 0.9967 |
| PG 18:1_20:3 | -0.0615 | 13.8908 | 0.0317 | 0.0005 | 0.0005 | 0.1592 | 0.0094 | 0.0014 | 0.0039 | 0.1569 | 0.6686 | 0.9231 | 0.3622 | 0.9873 | 0.9745 | 0.9967 |
| BMP 22:5_22:5 | 0.0612 | 13.8142 | 0.0012 | 0.2013 | 0.3596 | 0.0012 | 0.0711 | 0.0095 | 0.0001 | 0.4864 | 0.8319 | 0.6634 | 0.9013 | 0.5994 | 0.9745 | 0.9972 |
| PG 18:2_20:4 | -0.0556 | 12.5645 | 0.0262 | 0.0017 | 0.0995 | 0.1701 | 0.0746 | 0.0594 | 0.0032 | 0.0720 | 0.3655 | 0.1743 | 0.3307 | 0.7354 | 0.9745 | 0.9967 |
| BMP 18:0_22:5 | -0.0432 | 9.7590 | 0.1017 | 0.0000 | 0.0186 | 0.2283 | 0.1339 | 0.0129 | 0.1197 | 0.0025 | 0.0790 | 0.1253 | 0.0235 | 0.2323 | 0.7370 | 0.7578 |
| PG 18:3_22:6 | -0.0116 | 2.6136 | 0.0057 | 0.0201 | 0.0156 | 0.0250 | 0.0841 | 0.0477 | 0.0138 | 0.8895 | 0.9117 | 0.3840 | 0.4590 | 0.8161 | 0.9745 | 0.9967 |
| PG 22:4_22:6 | -0.0032 | 0.7160 | 0.0110 | 0.0251 | 0.1346 | 0.2705 | 0.1861 | 0.0485 | 0.0003 | 0.4918 | 0.4876 | 0.0564 | 0.1164 | 0.4217 | 0.9745 | 0.9972 |
| BMP 22:4_22:6 | 0.0024 | 0.5398 | 0.0649 | 0.1576 | 0.2978 | 0.0666 | 0.0271 | 0.0440 | 0.0051 | 0.1013 | 0.8654 | 0.9107 | 0.4026 | 0.6800 | 0.9213 | 0.9967 |

B

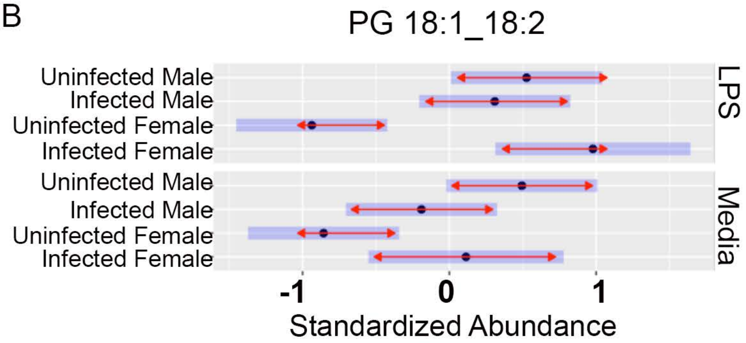


**Supplementary Figure 2. Elastic Net Machine Learning Model identifies a *S. mansoni* induced sex-specific lipidomic profile.** (A) Table of the 20 lipid features retained from an elastic net machine learning model. Negative values of the coefficient indicate that an increase in abundance of the compound is on average associated with a decrease in the probability that the sample comes from a female animal. (B) Visualization of the top lipid feature showing sex related differences in groups following LPS stimulation.
